# Microbiome- and Host Inflammasome-Targeting Inhibitor Nanoliogmers are Therapeutic in Murine Colitis Model

**DOI:** 10.1101/2024.02.20.581256

**Authors:** Sadhana Sharma, Vincenzo S. Gilberto, Cassandra L. Levens, Anushree Chatterjee, Kristine A. Kuhn, Prashant Nagpal

**Affiliations:** Sachi Bio, 685 S Arthur Avenue, Colorado Technology Center, Louisville, CO 8002, USA; Division of Rheumatology, Department of Medicine, University of Colorado Anschutz Medical Campus, Aurora, CO 80045, USA

**Keywords:** Autoimmune disease, Inflammatory Bowel Disease (IBD), Crohn’s Disease (CD), Ulcerative Colitis (UC), Inflammasome-inhibitor, Microbiome-targeting therapy, Nanoligomer

## Abstract

Autoimmune and autoinflammatory diseases account for more than 80 chronic conditions affecting more than 24 million people in the US. Amongst these autoinflammatory diseases, non-infectious chronic inflammation of the gastrointestinal (GI) tract causes inflammatory bowel diseases (IBD), primarily Crohn’s and Ulcerative Colitis (UC). IBD is a complex disease, and one hypothesis is that these are either caused or worsened by compounds produced by bacteria in the gut. While traditional approaches have focused on pan immunosuppressive techniques (e.g., steroids), low remission rates, prolonged illnesses, and increased frequency of surgical procedures have prompted the search for more targeted and precision therapeutic approaches. IBD is a complex disease resulting from both genetic and environmental factors, but several recent studies have highlighted the potential pivotal contribution of gut microbiota dysbiosis. Gut microbiota are known to modulate the immune status of the gut by producing metabolites that are encoded in biosynthetic gene clusters (BGCs) of the bacterial genome. Here we show, a targeted and high-throughput screening of more than 90 biosynthetic genes in 41 gut anaerobes, through down selection using available bioinformatics tools, targeted gene manipulation in these genetically intractable organisms using Nanoligomer platform, and identification and synthesis of top microbiome-targets as a Nanoligomer BGC cocktail (SB_BGC_CK1, abbreviated as CK1) as a feasible precision therapeutics approach. Further, we used a host-directed immune-target screening to identify NF-κB and NLRP3 cocktail SB_NI_112 (or NI112 for short) as a targeted inflammasome inhibitor. We used these top two microbe- and host-targeted Nanoligomer cocktails in acute and chronic dextran sulfate sodium (DSS) mouse colitis and in TNF^ΔARE/+^ transgenic mice that develop spontaneous Crohn’s like ilitis. The mouse microbiome was humanized to replicate that in human IBD through antibiotic treatment followed by mixed fecal gavage from 10 human donors and spiked with IBD-inducing microbial species. Following colonization, colitis was induced in mice using one week of 3% DSS (acute), or six weeks of 3 rounds of 2.5% DSS induction for a week followed by one week of no DSS (chronic colitis model). Both Nanoligomer cocktails (CK1 and NI112) showed a strong reduction in disease severity, significant improvement in disease histopathology, and profound downregulation of disease biomarkers in colon tissue as assessed by multiplexed ELISA. Further, we used two different formulations of intraperitoneal injections (IP) and Nanoligomer pills in the chronic DSS colitis model. Although both formulations were highly effective, the oral pill formulation demonstrated a greater reduction in biochemical markers compared to IP. A similar therapeutic effect was observed in the TNF^ΔARE/+^ model. Overall, these results point to the potential for further development and testing of these inflammasome-targeting host-directed therapy (NI112), and more personalized microbiome-cocktails (CK1) for patients with recalcitrant IBD.

TOC GRAPHIC

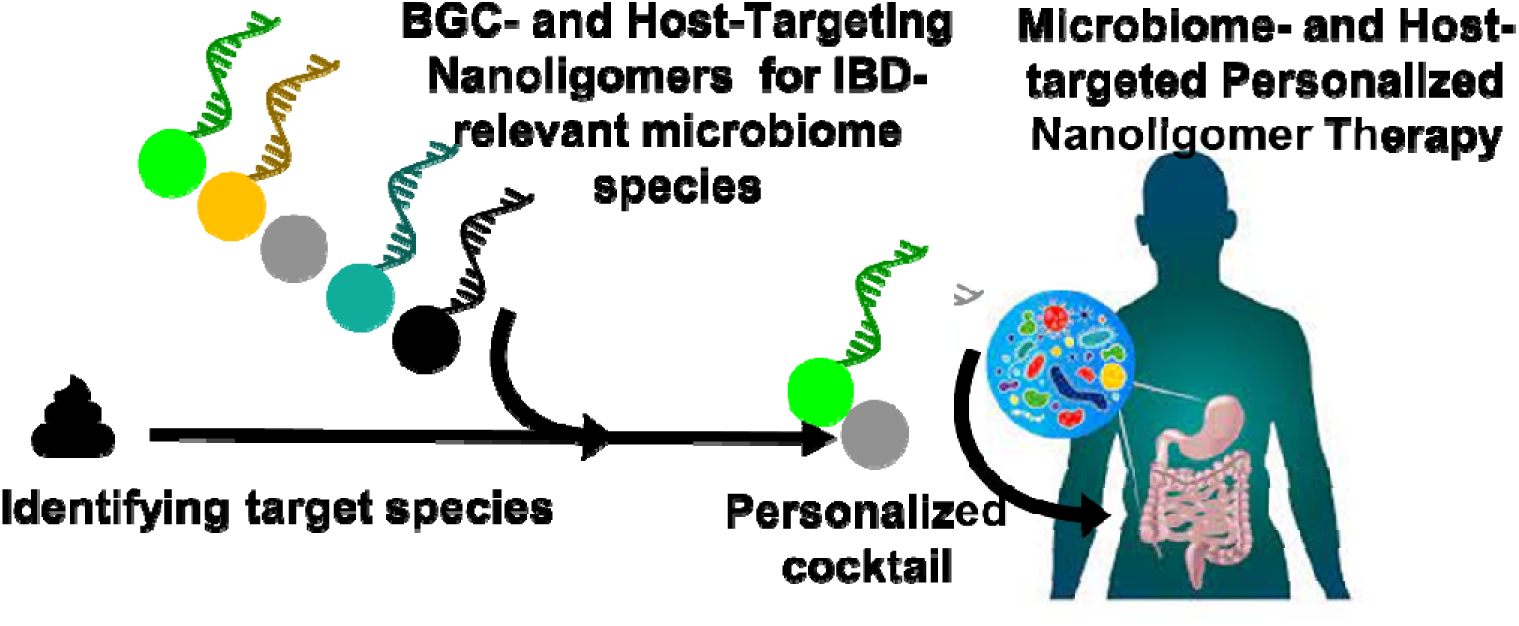

## INTRODUCTION

### Inflammatory Bowel Disease (IBD) is a Complex Illness with Inadequate Treatment Options

Approximately 1.6 million Americans and millions worldwide suffer from Crohn’s disease (CD) or Ulcerative colitis (UC), collectively described as IBD.^1,2^ The etiology of IBD is not entirely understood, but broad evidence indicates that the hallmark chronic inflammation stems from abnormal immune responses and alterations in gut microbiota, triggered by diet or other environmental causes in genetically susceptible individuals.^3–19^ The standard of care for IBD varies depending on disease severity and other factors, but it typically includes immunosuppressants like steroids that increase the long-term risk of colorectal cancer and achieve complete remission in less than half of patients.^20^ Given the immunosuppressants are not adequate for all patients and low remission rates, researchers have turned to more targeted immunotherapies and altered gut microbiomes for precision and personalized therapies.

### Altered Microbiome is a Target of Novel Therapeutics for IBD

Delivery of selected microbes (bugs as drugs),^21^ fecal matter transplants (FMT), and genetically engineered probiotics have been proposed as potential therapies targeting the microbiome. Each approach is designed to shift the composition and relative abundance of bacteria to better match individuals without IBD, with the hope that such a shift will restore an anti-inflammatory environment and eliminate the bacterial metabolites that contribute to the altered immune response. However, discouraging results of clinical trials^22–26^ and multiple reports of adverse events including the risk of disease transmission and morbidity^27^ may ultimately limit further development. More recently, interest in shaping the microbiome has focused on the possibility of altering the milieu of pro- and anti-inflammatory metabolites through direct inhibition or activation of bacterial pathways involved in maintaining homeostasis in the intestine. The literature includes reports of dozens of metabolites that are altered by intestinal bacteria.^5–8,17–19,28–31^ However, translating this information into viable therapeutic approaches has been challenging because the process of targeting and altering specific genes or gene products is slow, expensive, and often technically challenging in some bacterial species.^28,32^ In addition, the clinical utility of therapeutics developed with these methods may be limited by the diversity of microbiome disturbances observed across different patients with IBD. Thus, there is a need for a rapid, high-throughput, and cost-effective approach to modify the metabolite profile of the gut microbiome, ideally with the ability to tailor modifications to individual patients (personalized medicine), based on their gut microbiome profile.

### Host-Directed Targeted Immunotherapies for IBD

Several parts of the immune system are involved in disease initiation, disease development, and progression towards IBD pathophysiology. The misrecognition of healthy gastrointestinal (GI) tract cells (ileum in CD and colon in UC) is multitiered and typically initiated through CD4^+^ T-cell differentiation from T-helper (Th) Th1 and Th17 cells in CD, and Th2 cells in UC.^33–36^ This is followed by more upstream expression of pro-inflammatory cytokines like interferon-gamma (IFN-γ), IL-17/IL-22, and IL-13/IL-5 in CD and UC respectively. The immune cell infiltration is followed by the release of a cascade of downstream mediators of inflammation like tumor necrosis factor-alpha (TNF-α), IL-1β, and IL-6, that causes GI inflammation and tissue damage. The nuclear factor kappa-light-chain-enhancer of activated B cells (NF-κB) is a key inflammatory signaling regulator that is known to promote the expression of downstream pro-inflammatory genes, and strongly mediate the course of mucosal inflammation to amplify or accelerate IBD pathogenesis.^37,38^ Further, downstream activation of the NLRP3 (NOD-like receptor family, pyrin domain containing 3) inflammasome turns on the inflammatory cascade that leads to IBD pathology.^39^ Abnormal activation of this inflammasome, specifically NLRP3, has been actively investigated and associated with inflammation in colonic mucosa, epithelial cells, and immune cell infiltration.^40^ Therefore, inflammasome inhibition is being intensely investigated as a host-directed targeted therapy in autoimmune diseases, specifically IBD.^41,42^

### Nanoligomers are a Safe and Targeted Approach for Inflammation Mitigation

The remission rates for IBD treatment using generic anti-inflammatory therapies are low (< 50%),^20^ suggesting it may be necessary to target specific upstream mediators of inflammation (precision therapy), and/or mitigate or shape microbial metabolite profile depending on the patient-specific gut microbiota (personalized therapy) for therapeutic efficacy. In this context, Nanoligomers are a new therapeutic modality^43–48^ as gold nanoparticle-bound peptide nucleic acids (PNA) that can up- or down-regulate specific proteins with high specificity by binding to targeted DNA (transcriptional modulation) or mRNA (translational inhibition) (**Fig. 1A**).^43–48^ The nucleic acid-binding domain is a PNA, a synthetic DNA analog that demonstrates strong hybridization and specificity to their targets compared to naturally occurring RNA or DNA,^49^ and they exhibit no known enzymatic cleavage, leading to increased stability in human blood serum and mammalian cellular extracts.^50^ Given the low K_D_ (∼5 nM^45,46,49^), high binding specificity, minimal off-targeting,^45,46^ high safety profile, lack of any immunogenic response or accumulation in first pass organs, absence of any observable histological damage to organs even for long (>15-20 weeks) treatments,^45,46,48^ Nanoligomers can accelerate the development of effective and high-precision immune therapeutics through facile delivery to the target organs (gut here) using multiple routes of administration such as oral or injections. Nanoligomers are differentiated from the current standard of care (SOC) treatments in the following ways: 1) Reduce inflammation as evidenced by a decrease of multiple key cytokines targeted by different SOC and experimental therapies. 2) Expression of growth factors responsible for healing are kept at levels similar to, or higher than, healthy (or untreated) individuals. 3) Oral administration vs. current SOC that are mainly available as injectable or infusion.

**Fig. 1.**
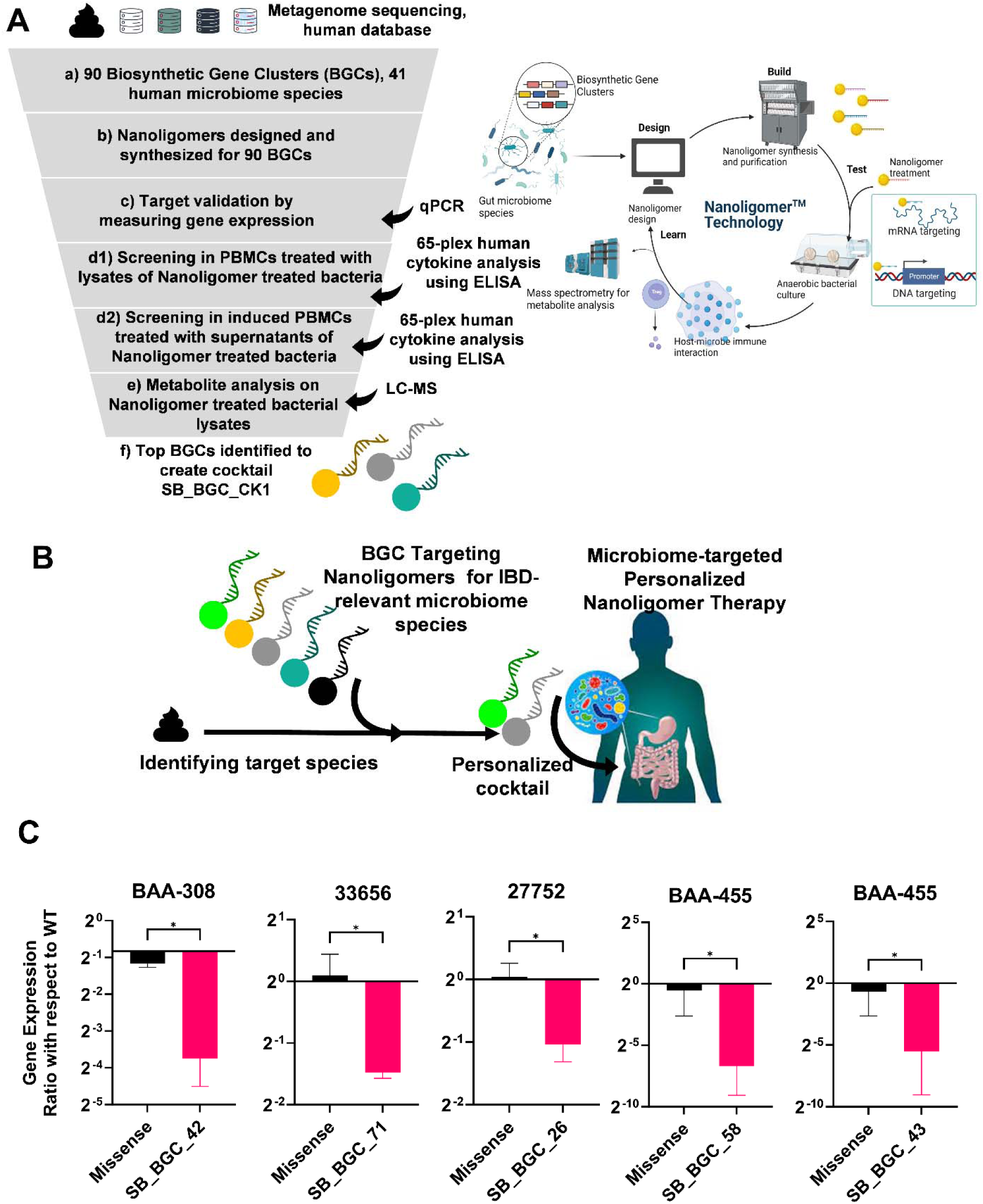
Nanoligomer platform for high-throughput screening of microbiome-targeted therapy in diverse and genetically intractable gut anaerobes. **A.** Schematic showing the proposed rapid and high-throughput screening methodology for down selection of target anaerobes and respective biosynthetic genes to modulate the gut metabolite profile and reduce gut inflammation. **B**. Utilizing the Nanoligomer-based BGC targeting approach shown in panel A, to create an individual-specific cocktail (based on their gut microbiome) and reduce inflammatory metabolites to mitigate gut inflammation, as a personalized IBD therapeutic approach. **C**. High-throughput gene targeting in relevant gut anaerobe species. The gene expression was measured using quantitative PCR (qPCR) and normalized with respect to WT anaerobes. As shown using respective missense molecules and BGC-targeting Nanoliogmers, the approach is highly effective (>2-4 fold downregulation of target genes) and targeted, minimizing potential off-targets. n=3 replicates for all measurements. \**P* < 0.05, \*\**P* < 0.01, and \*\*\**P* < 0.001, \*\*\*\**P* < 0.0001, Mean ± SEM, significance based on one-way ANOVA.

## RESULTS AND DISCUSSION

### In-vitro Screening and Testing in Donor-Derived Primary Human Peripheral Blood Mononuclear Cells (PBMCs)

Given the large number of gut anaerobes and their respective annotated biosynthetic genes or BGCs, we first selected 41 microbial species and 90 BGCs to be screened for IBD therapy. The down-selection was based on metagenomic sequencing of human fecal samples, and annotated BGCs and their metabolites from available databases such as antiSMASH 6.0^51^ and 5.0,^52^ BiG-FAM,^53^ DoBISCUIT^54^, and MIBiG 2.0^55^, using bioinformatics tool (**Fig. 1A**). Some examples of human microbiome species included *Bacteriodes fragilis, Eubacterium rectale, Faecalibacterium prausnitzii, Blautia coccoides, Blautia hansenii, Roseburia hominis, Clostridium sporogenes, Ruminococcus (Blautia) obeum, Akkermanisa muciniphila, Ruminococcus gnavus, Alistipes shahii* (complete list in **Fig. S1**). Some examples of BGCs within these species included C4Q21_RS06440, WP_014080837.1, Clospo_02864, ZP_01962381, WP_000945878.1. We prioritized the list of metabolites and strains that are hypothesized/known to produce strong immunomodulatory effects, and prioritized BGC targeting these metabolites, to assess their impact and prioritized them. Then we used our Nanoligomer™ platform to design and synthesize 90 Nanoligomers targeting genes or mRNA in 37 diverse and genetically intractable human gut anaerobes in a high-throughput, rapid process (**Fig. 1A**). Nanoligomers were designed to block ribosomal binding and prevent protein production.

To test and screen top microbiome-targeting Nanoligomers relevant for IBD, we used an *in vitro* gut-immune model (**Fig. 1B**). Microbiome cultures in their growth phase were treated with Nanoligomers targeting their respective BGCs. Given the high throughput nature of the screen and significant prior success in successful gene modulation of the targeted mRNA, ^43–48^ we spot-checked the gene expression modulation in these otherwise genetically intractable anaerobes. As shown in **Fig. 1C**, the designed Nanoligomers were able to selectively target the BGCs with high efficacy and were validated for modulation of gene expression of corresponding BGCs using qPCR. Overall, Nanoligomers targeting BGC met the minimum threshold of a 2-4 fold decrease (of target genes) in gene expression, with respect to both wild-type (WT) and missense (non-targeting) Nanoligomers. Following gene expression validation, the BGC modulated (using Nanoligomer treatment) and untreated cultures were used as: a) cell lysates (**Fig. 2A-E**, **3A**); or, b) cell supernatants (**Fig. 3B, C**), all filtered sterilized with 0.2 µm filter, and added to donor-derived human PBMCs, to monitor their effect on host immune system. The cell lysates could stimulate the PBMCs by themselves,^29,56^ whereas for cell supernatants, we used Phorbol 12-myristate 13-acetate PMA (25 ng/ml) and ionomycin (1 µg/ml) for 6 hours.^57^ All cytokines measured in the PBMC supernatants were analyzed using a multiplexed 65-panel ThermoFisher ProcartaPlex Human Immune monitoring panel (see Methods), and all treatments were compared with their respective untreated (WT) controls.

**Fig. 2.**
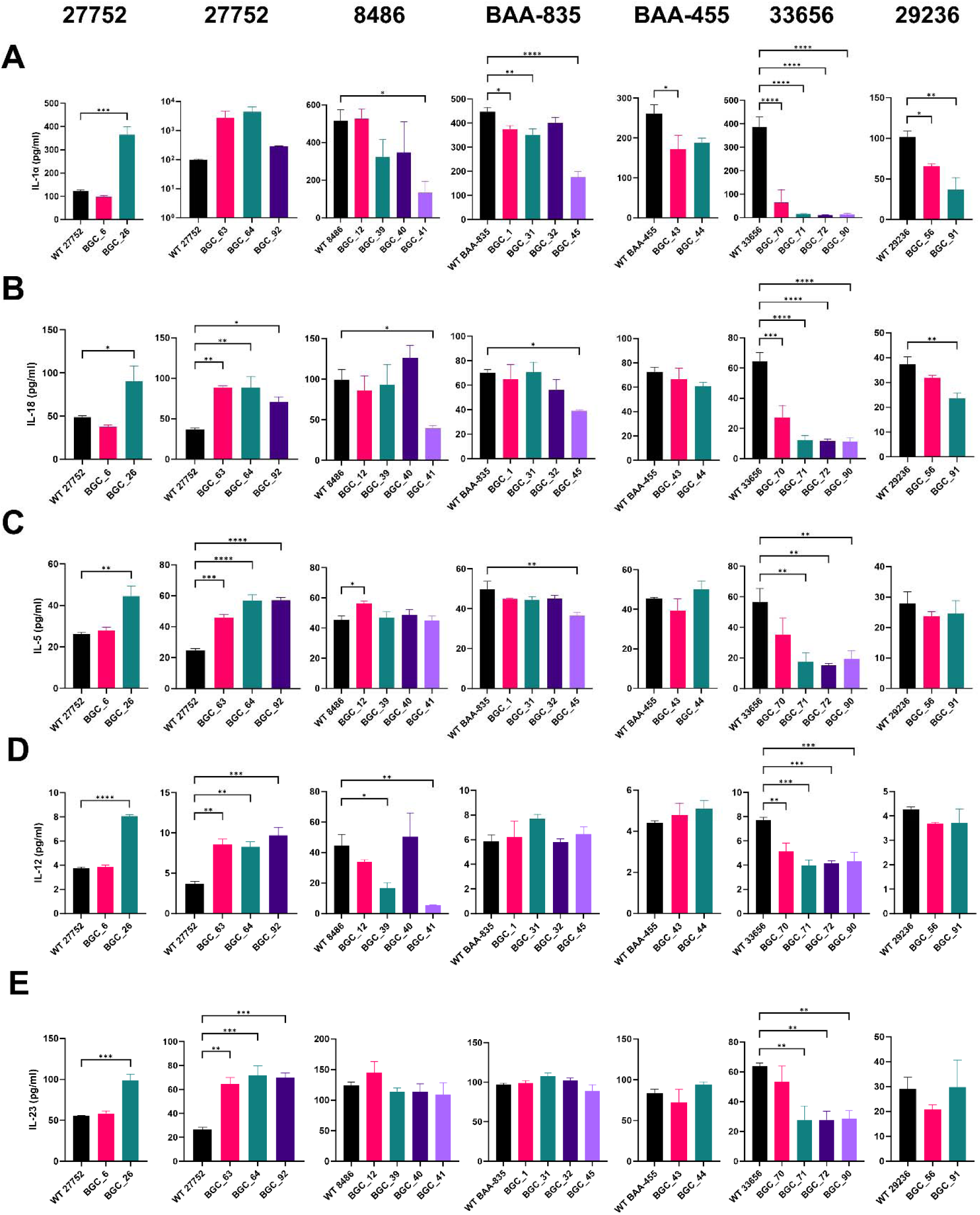
*In vitro* screening of microbiome species and gene targets using lysate treatments of donor-derived PBMCs. Different WT microbial species (*B. hansenii* (27752), *E. limosum* (8486), *A. muciniphila* (BAA-835), *P. xylanivorans* (BAA-455), *E. rectale* (33656), and *B. coccoides* (29236)) and their respective gene-downregulated cultures (using Nanoligomer treatments): 27752-BGCs 6, 26, 63, 64, and 92; 8486-BGCs 12, 39, 40, 41; BAA-835-BGCs 1, 31, 32, 45; BAA-455-BGCs 43, 44; 33656-BGCs 70, 71, 72, 90; and 29236-BGCs 56, 91, were lysed and used for 24-hour treatment of human PBMCs. Post-72-hour treatment with cell lysates, PBMC supernatants were analyzed for 65 cytokines and chemokines, to assess potential gut-immune interactions. To assess the treatments in this in vitro model for IBD therapy, the following disease-relevant cytokines are presented here: **A.** IL-1α, and **B**. IL-18 as inflammation-inducing pro-inflammatory cytokines. Additionally, upstream signaling cytokines **C**. IL-5, **D**. IL-12, and **E**. IL-23, were also measured for their role in T-cell differentiation for respective Th2 and Th1/Th17 inflammation pathways, for UC and CD respectively. \**P* < 0.05, \*\**P* < 0.01, and \*\*\**P* < 0.001, \*\*\*\**P* < 0.0001, Mean ± SEM, significance based on one-way ANOVA. N=3 for each group.

**Fig. 3.**
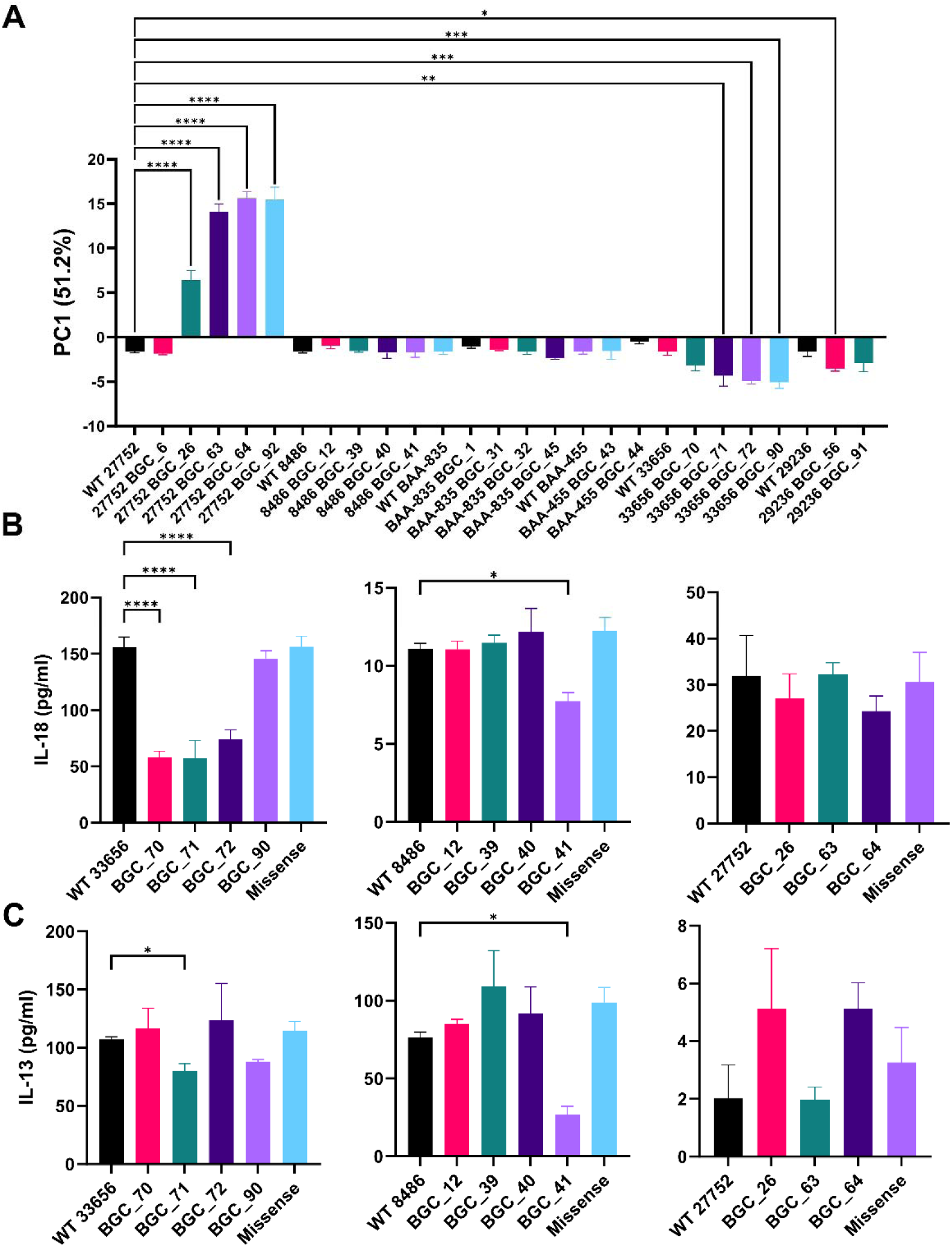
Ranking and validation of *in vitro* screening of microbiome-targets using the proposed gut-immune model, with both cell lysate and cell supernatant treatments. **A.** Principal component analysis used for combining the results of all 65-cytokines and chemokines, using the first principal component (PC1, 51.2%) from lysate treatment of PBMCs. The more negative values indicate a more anti-inflammatory response and more positive values show a more pro-inflammatory response. PC1 data was used for ranking microbiome-targeted therapeutics and probiotics. Cell supernatants from the top 3 strains (33656, 8486, and 27752), and their respective Nanoligomer-treated cultures were used for PBMC treatments (stimulated with PMA and Ionomycin for 6 hours). We used **B**. IL-18 as the key downstream pro-inflammatory cytokine, and **C**. IL-13 as more upstream signaling cytokine, to validate and advance top microbiome-targeted Nanoligomer therapeutics as cocktail CK1. \**P* < 0.05, \*\**P* < 0.01, and \*\*\**P* < 0.001, Mean ± SEM, significance based on one-way ANOVA. *n* = 3 for each group.

### Comparing immune expression profile of treated cell lysates

We screened different gut anaerobes (as potential probiotics) and their biosynthetic genes (as Nanoligomer targets) using this *in vitro* gut-immune model. To outline the key findings of this massive and high-throughput screening, we present 6 gut anaerobes as examples here: *Blautia hansenii* (ATCC 27752), *Eubacterium limosum* (ATCC 8486), *Akkermansia muciniphila* (BAA-835), *Pseudobutyrivibrio xylanivorans* (BAA-455), *Eubacterium rectale* (ATCC 33656), and *Blautia coccoides* (ATCC 29236). We chose 5 cytokines: IL-1α (**Fig. 2A**)^58^ and IL-18 (**Fig. 2B)** as downstream inflammation and IBD pathology mediators,); and IL-5 (**Fig. 2C**), IL-12 (**Fig. 2D**), and IL-23 (**Fig. 2E**) as more upstream signaling and T-cell differentiation cytokines, as outlined above. All 65-cytokine and chemokine results are shown in **Figs. S2-S8**. As seen here, except for BGC 6, 4 BGCs suppressed in *B. hansenii* (27752) lead to a strong pro-inflammatory response. This shows an overwhelming number of metabolites produced by *B. hansenii* that are strongly anti-inflammatory, which is why their omission/suppression leads to a strong pro-inflammatory response, compared to the WT. The remaining annotated BGC produced a small/weak immunomodulatory response (BGC 6), and hence this gut anaerobe could be a strong candidate for further testing as a probiotic for IBD therapy. Similarly, given the strong immunosuppression from BGCs 26,63, 64, and 92, these would be good candidates for a future immune upregulation therapeutic application (e.g. in oncology or infectious disease).

Next, we observed only one top anti-inflammatory BGC (showing a significant decrease in expression of IL-α or IL-18) can be easily identified from the remaining 4 gut anaerobes *E. limosum* (8486 BGC 41), *A. muciniphila* (BAA-835 BGC 45), *P. xylanivorans* (BAA-455 BGC 43), and *B. coccoides* (29236 BGC 92). Amongst these 4 strains and respective BGCs, only 2 strains *E. limosum* (8486) and *A. muciniphila* (BAA-835), and their respective BGCs 41 and 45 produce metabolites to influence both downstream immune suppression (**Fig. 2A, B**) and reduction of more upstream signaling cascade (BAA-835 BGC 45 in IL-5 **Fig. 2C**, 8486 BGC 41in IL-12 **Fig 2D**). IL-12, a cytokine with dual proinflammatory and immunoregulatory properties, plays a significant role in the pathogenesis of inflammatory bowel disease (IBD).^59^ Based on the above description of IL-5 in Th2 and IL-12 in Th1 cascades,^33–36^ 8486 BGC 41 would be a good Nanoligomer target for CD and BAA-835 BGC 45 Nanoligomer for UC therapy. This personalized microbiome-targeting therapeutic would be expected to be effective if *E. limosum was* found in abundance for CD patients, and *A. muciniphila* was found in the gut of UC patients, respectively. Several studies have noted the decreased abundance of mucin-degrading *A. muciniphila* in IBD patients and while it had been proposed as a strong candidate for therapeutic probiotics given possible regulation of host barrier function and immune response,^60^ its contradictory results in IBD have prevented further progress.^61^ Similarly, given strong butyrate, acetate, propionate, and lactate production by *E. limosum*, it has been proposed as a probiotic for IBD.^62^ We believe the BGC results presented here have shed more light on the apparent contradiction, and open new avenues for the use of Nanoligomer BAA-835 BGC 45 with *A. muciniphila* and 8486 BGC 41 with *E. limosum* probiotic, to counter the specific adverse impact of their respective biosynthetic gene metabolites in IBD therapy.

Another controversial gut anaerobe we picked in this study was *E. rectale* (33656). Given the high butyrate production and the possibility of promoting good colon health,^63^ potential implications of *E. rectale* as an IBD organism and its role in inducing colorectal cancer through NF-κB activating metabolites have been controversial.^64^ Our results here show that a large majority of *E. rectale* BGC inhibition (70, 71, 72, 90) are strongly anti-inflammatory (significant downregulation of IL-23), indicating they produce strongly pro-inflammatory metabolites (**Fig. 2E**). IL-23, a proinflammatory cytokine, plays a crucial role in IBD, and is implicated in driving inflammation and tissue damage in the gastrointestinal tract. Specifically, IL-23 promotes the differentiation and activation of T helper 17 (Th17) cells, which are known to be involved in the pathogenesis of IBD.^59^ Given the large magnitude of reduction in all downstream inflammatory cytokines (**Fig. 2A, B**), and upstream signaling molecules (**Fig. 2C, D, E**) related to CD and UC, these Nanoligomers, especially 33656 BGC71 and 72, were selected as cocktail lead molecules for *in vivo* IBD testing.

Further screening analysis using all 65-cytokines (**Figs. S2-S8**) using Principal Component Analysis (PC1, 51.2%). More negative values of PC1 correspond to a more anti-inflammatory response, whereas more positive values correspond to a more pro-inflammatory response.^43,44^ WT value is shown as a reference point. The analysis showed the same results with three *E. rectale* BGCs (71, 72, 90) identified as top targets (**Fig. 3A**). Additionally, we utilized these PC1 screening and ranking for all 41 strains and 90 BGCs to identify the following strains and their top respective BGCs as IBD therapeutic cocktail SB_BGC_CK1 (or CK1 for short): *E. rectale* (33656), *B. coccoides* (29236), *A. muciniphila* (BAA-835), and *A. shahii* (BAA-1179).

### Comparing immune expression profile of treated cell supernatants

We showed the effect of cell-lysates (above), and how top Nanoligomer targeting specific BGCs were identified. We also treated donor-derived PBMCs using cell supernatants of the untreated and top Nanoligomer-treated bacterial cultures. While the 65-cytokine immune profiles are different, key outcomes and results were consistent. One key difference was the IL-18 expression was the key pro-inflammatory output in cell supernatants (**Fig. 3B**), not IL-1α, IL-1β, IL-17, etc., likely due to early profiling (6-hour supernatant treatment^57^ of PBMCs instead of 72-hour cell-lysate treatment protocol^29,56^ established in the gut-immune models used) of developing immune-modulation. Comparing the two models, top Nanoligomer molecules and targets were similar, although the ranking of *E. rectale* (33656) targets changed slightly. However, top BGC targeting Nanoligomers (SB_BGC_71, 41 **Fig. 3B, C**) again showed significant 2-3 fold reduction of host inflammation (**Fig. 3B**) and upstream cytokine signaling (**Fig. 3C**). This was also verified for key Nanoligomer molecules selected for *in vivo* evaluation in IBD mouse models.

### Nanoligomers alter metabolite profiles of genetically intractable gut anaerobes

To assess whether the changes in metabolite profile can modulate immune responses, we collected lysates and supernatants from the Nanoligomer-treated microbiome and assessed the effect of BGC targeting on metabolites using LC-MS profiling. Here we show the data from the top 2 anti-inflammatory and top 2 pro-inflammatory response-inducing BGCs (33656 BGCs 71, 72, and 27752 BGCs 26, 63, **Figs. 2, 3A**). In analyzing the data, we just focused on the differential expression with respect to WT, and used p=0.05 as the significance screening threshold. In a differential expression of 33656 BGCs 71 and 72, more than 2000 targeted and untargeted metabolites were identified, and a significant fraction of them (>25%) were differentially expressed above the significance threshold (p<0.05) (65 and 66 metabolites from 200 targeted metabolites shown, **Fig. 4 A, B).** While differential expression metabolite profile for 33656 BGCs was very similar, except for sphingolipid biosynthesis metabolite sphinganine 1-phosphate (>2-fold enhanced compared to WT) in BGC 72, similar analysis of 22752 BGCs showed significantly lower number of differentially expressed metabolites (∼5-10%) with significant differences between the two BGCs (27752 BGCs 26, 63, **Fig. 4 C, D)**. The fold-change observed for 27752 BGCs was also lower, in BGC 63, compared to 26. Another important insight into the role of these metabolites for IBD therapy comes from short-chain fatty acid (SCFA) derivatives in 33656, which are important for colonocyte growth and metabolism and promoting a healthy gut microbiome, as well as some selected metabolites in the tryptophan-indole pathway. SCFAs participate in the maintenance of intestinal mucosa integrity,^65^ improve glucose and lipid metabolism,^66^ control energy expenditure, and regulate the immune system and inflammatory responses.^67^ Given their strong anti-inflammatory role,^68^ modulation of gut epithelial cells^69^ and preventing/reversing gut-dysbiosis linked to IBD and colorectal cancer,^70^ and promoting a healthy gut microbiome,^71,72^ the more than 2-fold upregulation seen with top BGC targets further supports their testing in an *in vivo* IBD model. Additionally, using the indole and tryptophan metabolite analysis, we clearly see significant upregulation of these key anti-inflammatory and barrier integrity strengthening metabolites by more than 2-fold (**Fig. 4A, B**).

**Fig. 4.**
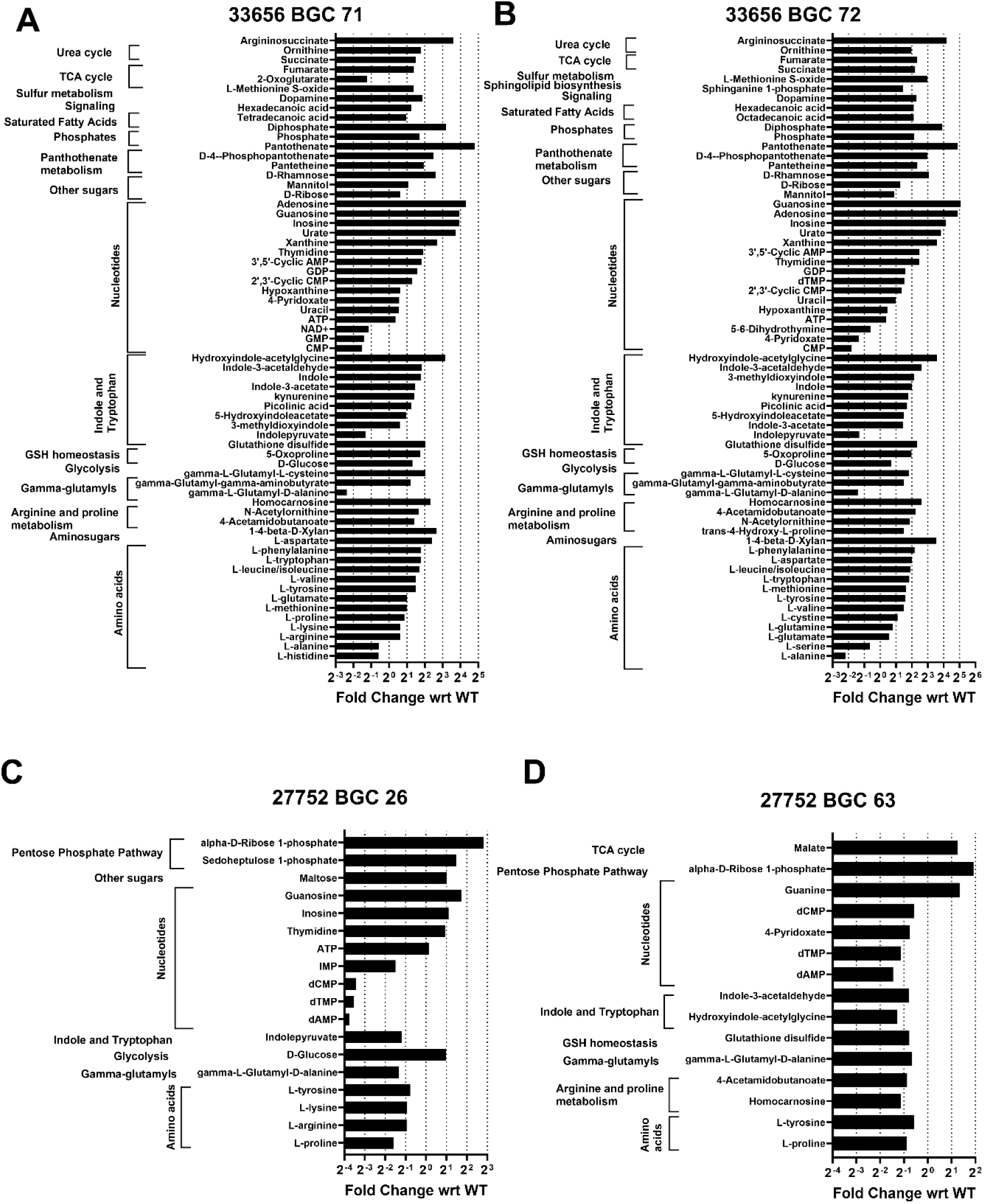
Metabolite analysis for top 2 pro- and anti-inflammatory microbiome treatments. Differential metabolite expression analysis showing fold-change between cell lysates of: **A.** WT 33656 vs BGC 71-suppressed 33656. **B.** WT 33656 vs BGC 72-suppressed 33656. **C**. WT 27752 vs. BGC 26-suppressed 27752; and **D**. WT 27752 vs. BGC 26-suppressed 27752. A significant threshold of p<0.05 was used. *n* = 3 for each group.

### Nanoligomers are Therapeutic in both acute and chronic *in vivo* colitis (DSS) mouse models

Following in vitro screening, ranking, and metabolite validation of the top two microbiome- and host-targeted Nanoligomer cocktails CK1 and NI112, we assessed the two lead Nanoligomers in the Dextran Sodium Sulfate (DSS) mouse colitis model. First, we treated WT C57BL/6 mice with antibiotics (added to drinking water) for one week followed by oral gavage with a mixed fecal slurry from 10 IBD patients spiked with 4 gut anaerobe species identified above: *E. rectale* (33656), *B. coccoides* (29236), *A. muciniphila* (BAA-835), and *A. shahii* (BAA-1179). Following one week of stable colonization, we treated mice with DSS in the drinking water. For acute 1-week treatment, we used 3% DSS water, and for chronic DSS model, we used 3 cycles of 5 days of 2.5% DSS treatment, followed by 5 days of normal drinking water (for recovery). The disease activity index (DAI) scores were monitored during the experiment, and after 1-week (for acute) and 3-weeks (for chronic) studies, the mice were euthanized, their colons extracted for multiplexed tissue ELISA to monitor inflammation, and some samples were used to conduct histological evaluation. Non-DSS treated and non-fecal gavaged mice were used as negative/healthy controls (shown as Non-DSS), and mice with sham/vehicle treatment (sterile saline in a 1-week study and enteric-coated pills with filler alone in a 3-week study) were used as diseased positive control mice, shown as DSS + Sham. As shown here, both lead molecules (NI112 and CK1) showed a rapid reduction in the disease activity index (DAI) scores with just 1-week of treatment (**Fig. 5A**). Mice with sham treatment showed high disease severity, characterized by loose/watery stool and blood. However, the mice with NI112 and CK1 treatment showed complete prevention of disease in most mice within a week. This difference in colitis pathology was supported by histology samples with sham-treated mice showing significantly swollen colons with immune infiltration shown with marked arrows, and the two treated groups showing healthy colons with no or minimal immune cell infiltration (**Fig. 5B**). Further biochemical assessment of homogenized colon tissue (see Methods for Multiplexed ELISA) revealed a significant reduction in inflammation, correlating with DAI scores and histology. Using downstream inflammation-inducing and upstream signaling cytokines and chemokines as biomarkers of inflammation and resulting pathophysiology, we observed a significant reduction in inflammation. For example, using TNF-α as one downstream biomarker, we observed a 90% reduction with NI112 and a 66% reduction with CK1 compared with positive control or sham-treated diseased mice. The values were reduced back to the level of healthy mice, and statistically identical negative control, as shown in **Fig. 6**. Similar trends were seen with upstream biomarkers like IL-12 (reduction of 84% with NI112 and 70% with CK1), IL-23 (reduced by 95% with NI112 and 85% with CK1), IL-5 (97% reduced with NI112 and 89% with CK1), and IL-17 (85% reduced with NI112 and 50% with CK1), using IP route of administration as shown in **Fig. 6**. The only exception was M-CSF, which showed massive increase in protein expression compared to both negative and positive controls (>4-fold increase with NI112), indicating the potential upregulation of repair and wound healing via these growth factors.

**Fig. 5.**
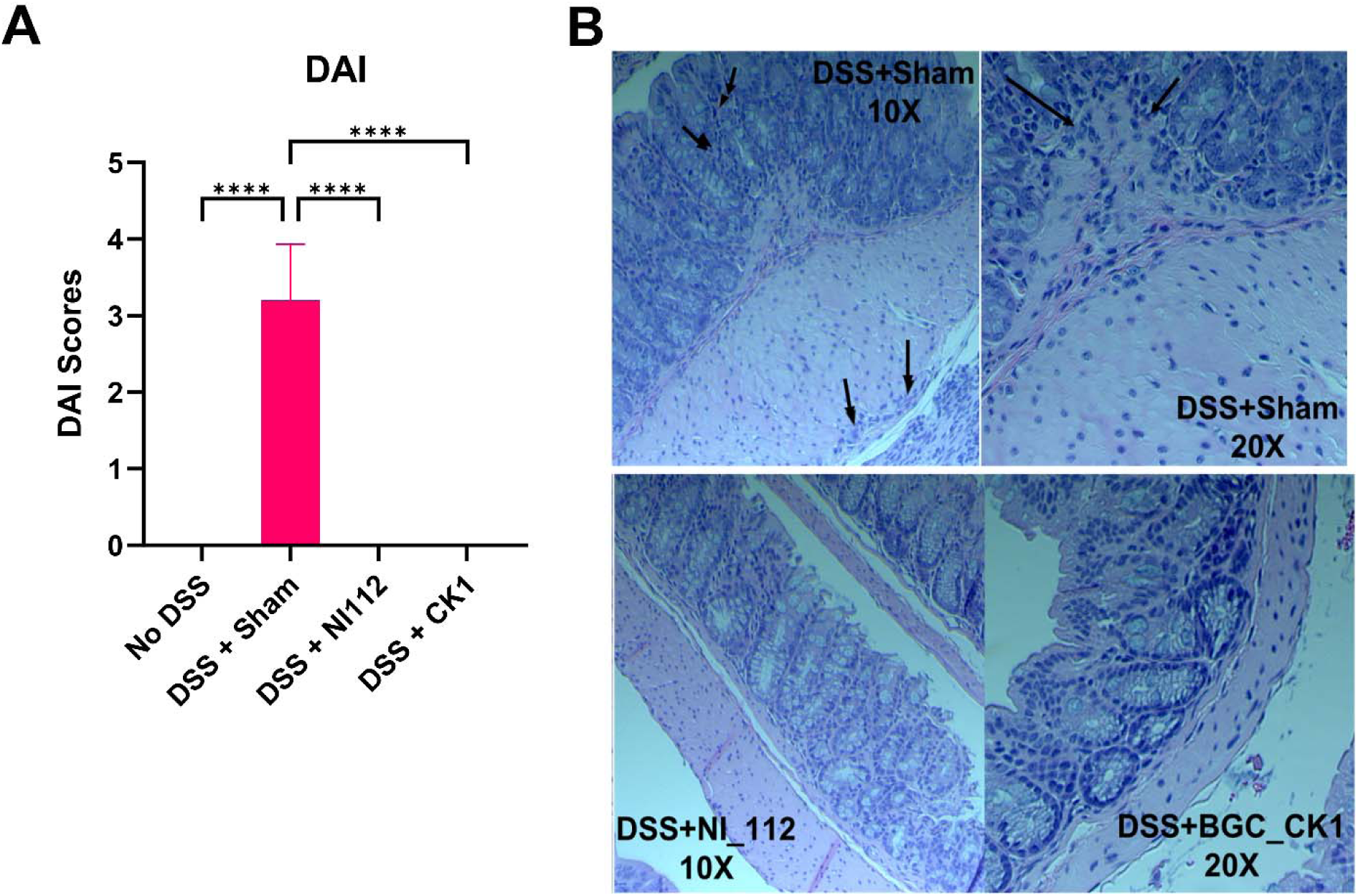
Disease severity and histological examination of microbiome-targeting (CK1) and host-targeting (NI112) Nanoligomer therapies in DSS-induced colitis mouse model. **A.** Disease severity, shown as disease activity index (DAI), is significantly reduced in both treatments (NI112 and CK1), at par with no DSS (or negative control) mice. The lack of loose stool or blood in stool for DSS-treated mice showed good clinical observation with lead Nanoligomers. **B.** Histologic examination using H&E staining showing significant swelling and immune cell infiltration in representative DSS-sham mice colon (top) and normal healthy colon with minimal or no-observable immune cell accumulation or colon shredding in NI112 and CK1 treated mice. The histology exams match the DAI scoring from the clinical evaluation of mice stool and health. \**P* < 0.05, \*\**P* < 0.01, and \*\*\**P* < 0.001, Mean ± SEM, significance based on one-way ANOVA. *n* = 5 for each group. Method of administration: intraperitoneal (IP) injection.

**Fig. 6.**
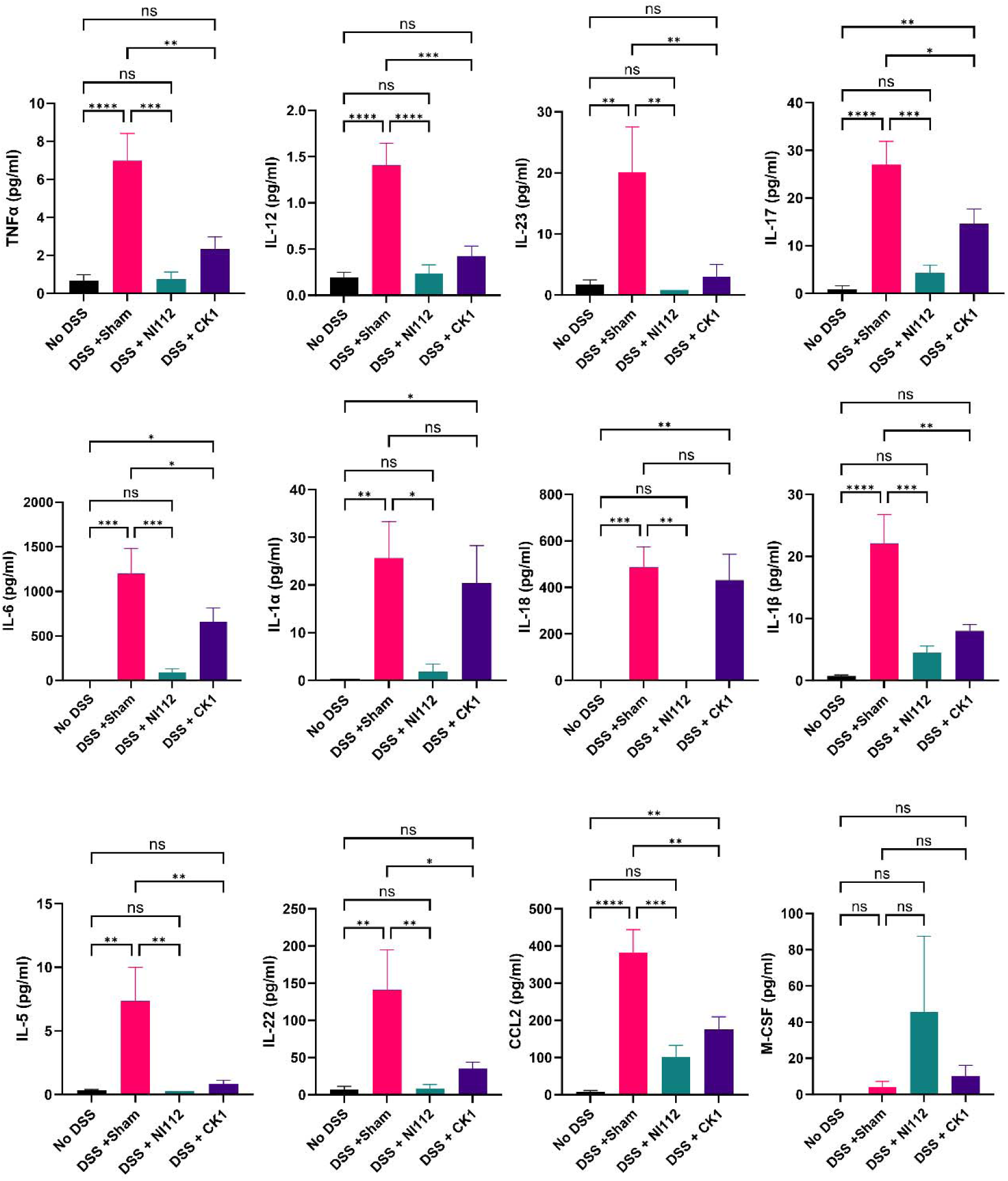
Biochemical assessment of mouse colon tissue ELISA reveals NI112 and CK1 Nanoligomer treatments ameliorate colitis biomarkers in an *in vivo* DSS mouse model. Significant reduction in 12-IBD relevant disease biomarkers was observed with NI112 and CK1 treatments in a 3% DSS mouse model after 1 week. This acute *in vivo* model demonstrates the ability to rapidly reduce disease severity and eliminate both upstream and downstream IBD biomarkers and cytokines participating in IBD pathophysiology, combining the mechanism of action of multiple therapies, and potentially providing immediate relief. The upregulation of growth factors like M-CSF shows potential for inducing wound healing/colon repair. \**P* < 0.05, \*\**P* < 0.01, and \*\*\**P* < 0.001, Mean ± SEM, significance based on one-way ANOVA. *n* = 5 for each group. Method of administration: intraperitoneal (IP) injection.

Due to the chronic recurrence of IBD disease and low remission rates, we further assessed the lead molecules in a 3-week chronic DSS IBD/colitis mouse model described above. We used two routes of administration here: IP injections and oral dosing using enteric-coated capsules (See Methods). Due to the difficulty in formulating the microbiome cocktail as oral capsules and similarity in responses observed between NI112 and CK1, we focused on NI112 therapy to test the efficacy in chronic colitis model and asses favorable route of administration for further translation (**Fig. 7**). Expectedly, the chronic DSS model also showed a massive reduction in pathology and showed decrease in key inflammatory markers in mice colon, with both routes of Nanoligomer treatments. The magnitude of reduction observed in inflammation biomarkers was more modest with NI112-IP treatment in these chronic DSS mice, as compared to sham-treated mice (reduction of ∼25-40% for most biomarkers, **Fig. 7**), indicating scope for improvement in therapeutic outcomes with dose escalation, dose frequency, and optimizing pharmacokinetic parameters with route of treatment. However, the NI112 treatment with the oral route (NI112-Pill) showed again massive reductions in inflammatory markers (>90% for most key cytokines, **Fig. 7**), at par or below healthy controls (No DSS mice). This indicates oral route may be a preferred method for therapeutic dosing and further pre-clinical and clinical testing of Nanoligomer lead molecules.

**Fig. 7.**
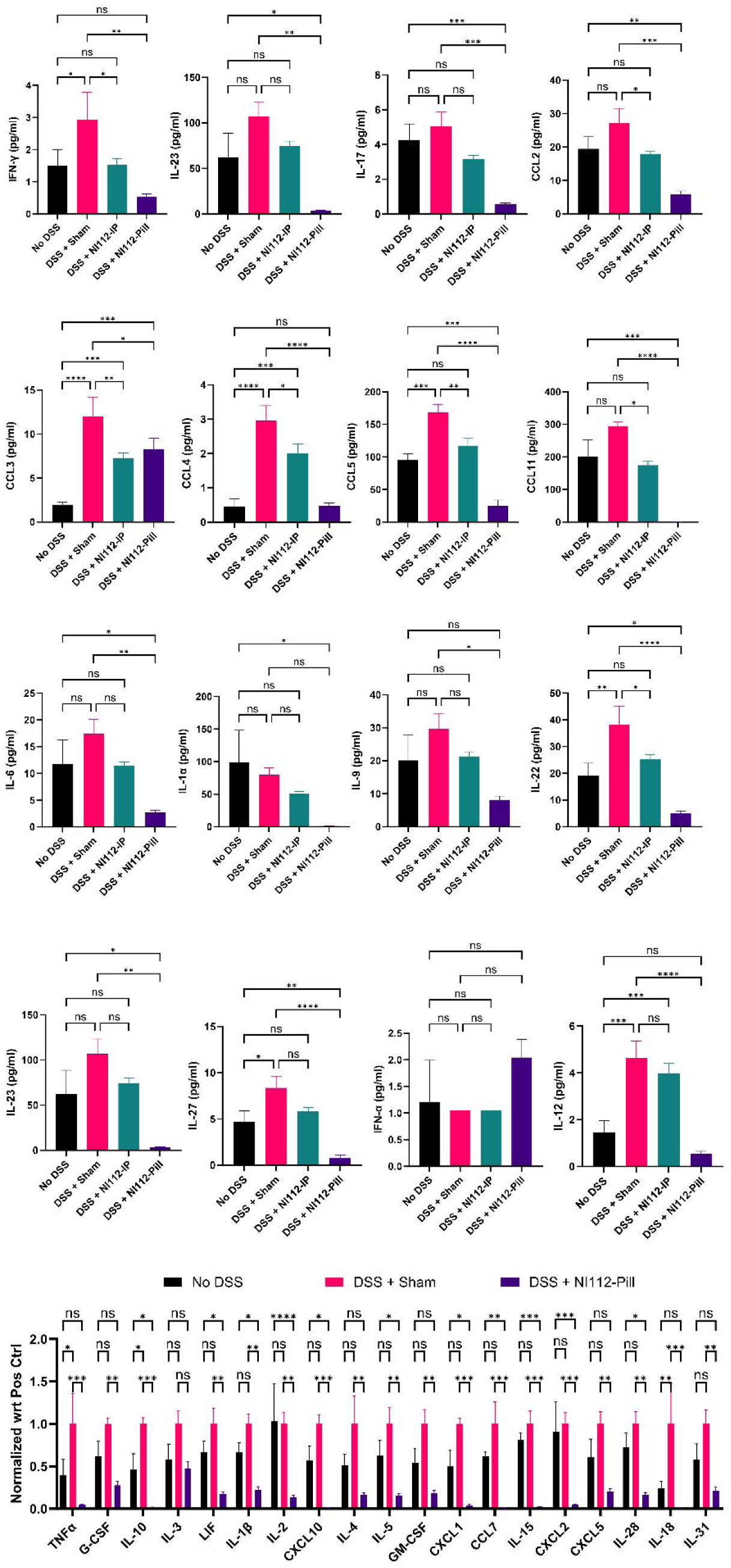
Nanoligomer treatments are therapeutic in chronic DSS mouse model. Multiple inflammatory and signaling cytokines and chemokines are significantly reduced using NI112 treatments in chronic DSS mouse model for IBD/colitis. While NII2 IP injections (DSS+ NI112-IP) 3x week 150 mg/kg dosing significantly reduces all IBD biomarkers in colon tissue, the impact of oral administration (DSS + NI112-Pill) was significantly higher, and restored all biomarkers to their healthy state (no DSS) or even lower. \**P* < 0.05, \*\**P* < 0.01, and \*\*\**P* < 0.001, Mean ± SEM, significance based on one-way ANOVA. *n* = 3-6 for each group (n=6 for No DSS and DSS+NI112-IP, n=3 for DSS+Sham, and n=5 for DSS+NI112-Pill). Methods of administration: intraperitoneal (IP) injection and oral (pills).

### Nanoligomer Therapeutic is Effective in TNF^ΔARE/+^ mouse model

Next, we tested the efficacy of lead Nanoligomer molecules in a genetically induced TNF^ΔARE/+^ mouse model. This model involves a mutation in the adenylate-uridylate-rich element (ARE) of the TNF gene, leading to increased stability of TNF mRNA and higher TNF production. Elevated TNF levels contribute to intestinal inflammation and colitis in these mice. TNF^ΔARE/+^ model is a standard model of human Crohn’s disease and is characterized by elevated TNF expression and inflammation in the ileum as is observed in Crohn’s disease patients.^73,74^ However, due to the focus of this study on murine colitis, we focused on mouse colon tissue analysis. Similar to the DSS model, the TNF^ΔARE/+^ mice were acclimatized for 2 weeks, and then treated with antibiotics for 5 days, and then recolonized with fecal gavage prepared from 10 different IBD patients (collected at the University of Colorado Anschutz). We also spiked the IBD fecal gavage with *E. rectale, A. muciniphila, Blautia coccoides,* and *Alistipes shahii* between weeks 6-7. Following one week of successful recolonization of the gut of these TNF^ΔARE/+^ with IBD relevant/representative microbiome, the mice were divided into three groups: 1) TNF^ΔARE/+^ + Sham (saline as sham, positive control), 2) TNF^ΔARE/+^+ NI112, and 3) TNF^+/+^ (negative control). After 4 weeks of treatment with 3 doses/week of 150 mg/kg NI112 administered IP, the mice were euthanized, and colon tissue samples were harvested for cytokine analysis. Evaluation of cytokines validated the efficacy of Nanoligomer treatment, as shown by the significant reduction of key IBD-associated inflammatory cytokines including TNF-α, IL-23, IL-17A, IL-12p70 (**Fig. 8**). While TNF^ΔARE/+^ mice show lower inflammation compared to DSS-induction, the biomarkers were reduced to their respective negative controls. These results provide strong evidence for the efficacy of lead molecules in multiple *in vivo* IBD/colitis mouse models.

**Fig. 8.**
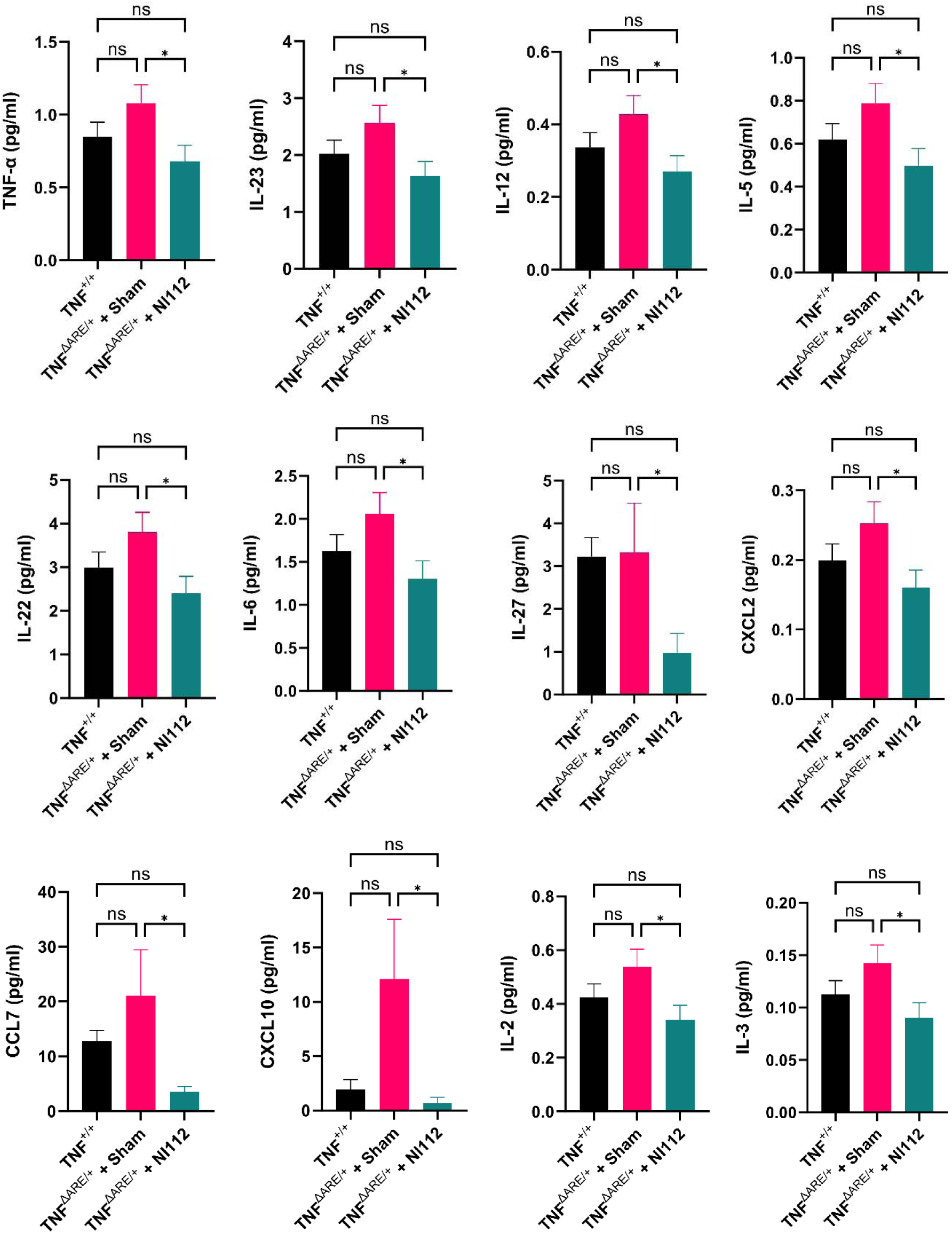
Genetic TNF^ΔARE/+^ mouse model confirms effectiveness of Nanoligomer therapeutic approach for IBD. 12 IBD inflammatory biomarkers in mouse colon tissue show effectiveness of NI112 Nanoligomer therapy in genetic TNF^ΔARE/+^ mouse model. TNF-negative mice were used as negative control (Neg Ctrl), saline (sham-treated) TNF^ΔARE/+^ mice were used as positive control (Pos Ctrl), and NI112-dosed (3x week, 150 mg/kg) TNF^ΔARE/+^ mice were used as treatment group (NI112 Trt). \**P* < 0.05, \*\**P* < 0.01, and \*\*\**P* < 0.001, Mean ± SEM, significance based on one-way ANOVA. *n* = 3-6 for each group (n=6 for NI112 Trt, n=3 for Neg Ctrl, and n=5 for Pos Ctrl). Method of administration: intraperitoneal (IP) injection.

## CONCLUSIONS

In conclusion, our study demonstrates screening of 41 microbiome species and 92 biosynthetic genes led to the identification of the top microbiome-targeting Nanoligomer cocktail CK1 for reducing inflammatory metabolites in the gut of IBD patients. This microbiome-targeting Nanoligomer cocktail CK1, as well as a host-targeting NF-κB and NLRP3 cocktail NI112, were screened in two different *in vivo* mouse models of intestinal inflammation: DSS-induced colitis and transgenic TNF^ΔARE/+^ mice. The Nanoligomer cocktails effectively mitigated inflammation in both models, and rescued mice by reducing disease severity, restoring all disease biomarkers to their healthy state, minimizing inflammation, and exhibiting minimal to no immune cell infiltration. These outcomes were comparable to negative control mice. We employed two different formulations for NI112: IP injections and enteric-coated oral capsules. While both methods showed strong efficacy, oral dosing far surpassed the efficacy seen with IP injections (as measured using colon tissue ELISA) and would be the preferred method for further translation. The study limitations include smaller group sizes for longer studies, and the use of NI112 only for oral dosing (due to formulation challenges with CK1). These findings suggest the potential for further development of these Nanoligomer therapies as personalized microbiome-targeting therapy (CK1, depending on each patient’s individual microbiome profile) for patients unable to achieve remission, as well as precision inflammasome-targeting host-directed therapy, with the potential to improve remission rates and reduce painful surgical procedures for chronic and recurring IBD in patients, that can benefit individuals affected by such debilitating autoimmune diseases like IBD.

## MATERIALS AND METHODS

### Bacterial strains and culture conditions

All the strains used in this study were obtained either from the American Type Culture Collection (ATCC) (Rockville, MD, USA) or Leibniz Institute DSMZ (Germany). All strains, except E. coli ATCC 10798 (aerobic culture), were cultured anaerobically (20% CO2) at 37°C in the broth culture medium specified in Figure S1. Ten different broth media were used. Mega Media^75^, PYAG (Peptone-yeast extract (PY) broth supplemented with 1% glucose and 33 mM sodium acetate) medium,^76^ YCFAG (yeast extract, casitone, fatty acid and glucose) medium,^77^ ATCC® Medium 1249: Modified Baar’s medium for sulfate reducers, ATCC® Medium 1827, ATCC® Medium 2971: GS2+ Cellobiose, and ATCC® Medium 2167: Haemophilus Test Medium were prepared as per the reference. Peptone-Yeast extract supplemented with 1% glucose (PYG AS-822), Chopped Meat broth (AS-811), and Chopped Meat Medium w/ Carbohydrates (AS-823) were procured from Anaerobe Systems, Morgan Hill, CA. *Coprococcus catus* ATCC 27761 and *Eubacterium eligens* ATCC 27750 were recovered from freeze-dried powder on Tryptic Soy Agar with 5% Sheep Blood (defibrinated) plates (ATCC medium 260) and then cultured in respective media.

### Nanoligomer Design and Synthesis

Nanoligomers were specifically designed and produced by Sachi Bio Inc., as described in detail elsewhere. ^43–47^ These Nanoligomers are comprised of a nanobiohybrid molecule comprised of an antisense peptide nucleic acid (PNA) moiety ^43,44,47^ conjugated to a nanoparticle^43,44,47^ for enabling enhanced delivery and cellular membrane penetration. The PNA segment of the Nanoligomer was synthesized on a Vantage peptide synthesizer (AAPPTec, LLC) employing solid-phase Fmoc chemistry. Fmoc-PNA monomers (A, C, and G monomers shielded with Bhoc groups) were purchased from PolyOrg Inc. Post-synthesis, the PNAs were linked with gold nanoparticles and subsequently purified using size-exclusion filtration. The conjugation process and the concentration of the refined solution were monitored by measuring absorbance at 260 nm (for PNA detection) and 400 nm (for nanoparticle quantification).

### Nanoligomer treatment, bacterial supernatant collection, and lysate preparation

All the Nanoligomer treatment experiments were performed in a 96-well plate. Anaerobic bacterial cells were treated with the relevant Nanoligomer (10 µM in MilliQ water) designed against specific BGC or missense Nanoligomer (10 µM in MilliQ water) for 21 hours (n= 3). A volume of MilliQ water equal to that of the Nanoligomer was added to the control. After treatment, bacterial supernatants were collected and stored at −80°C for future use. For lysate preparation, cells were centrifuged, washed, suspended in PBS, and heat-inactivated at 65°C for 30 min. The lysates were aliquoted and stored at −80°C for future use.

### RNA extraction and real-time quantitative polymerase chain reaction (RT-qPCR)

RNA was extracted using a miRNeasy Micro Kit (Qiagen, USA) per manufacturer protocol. Genomic DNA was removed using the TURBO DNA-free™ Kit. Zymo RNA Clean & Concentrator™-5 was used to inactivate the TURBO™ DNase. The resultant RNA (40Lng) was reverse transcribed to cDNA using a High-Capacity cDNA Reverse Transcription Kit (Applied Biosystems) using custom gene-specific reverse primers (Table 1). qPCR was performed using Fast SYBR Green Master Mix (Applied Biosystems) and custom primers (Integrated DNA Technologies, Inc., Table S1) on Bio-Rad CFX96 Touch Real-Time PCR Detection System. Gene expression for wcfR was analyzed following the delta-delta Ct method relative to the reference gene, DNA-directed RNA polymerase subunit beta, rpoc.

**Table 1.**
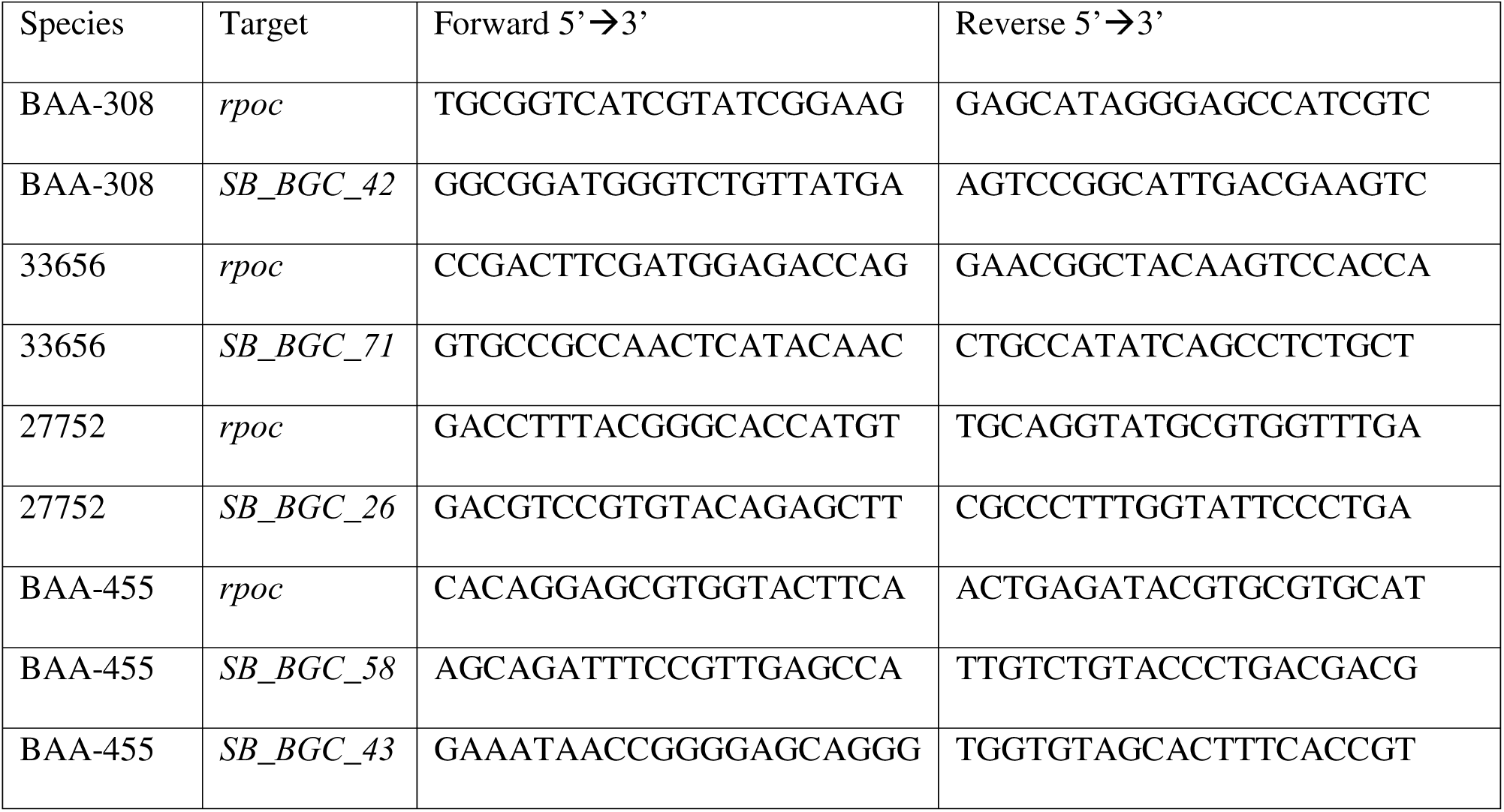
Primers used for qPCR.

### PBMC stimulation with bacterial supernatant or lysate

Human PBMCs used in this study were purchased from Zen Bio, Inc (Research Triangle, NC) and stored in the vapor phase of liquid nitrogen. On the day of the experiment, cells were thawed as per the manufacturer’s instructions and suspended in complete medium (RPMI-1640 w/ Glutamax + 10% heat-inactivated fetal bovine serum (FBS) + 1% Penicillin /Streptomycin. 10 µL of the bacterial supernatant (or respective blank medium as no treatment control) or lysate (or PBS as no treatment control) was added to 500k PBMCs (total volume 200 µL). For supernatant-based PBMC stimulation, a stimulating agent cocktail supplemented with Phorbol 12-myristate 13-acetate (PMA) and Ionomycin at an effective concentration of 25 ng/mL and 1 μg/mL, respectively was also added. The cell culture supernatants were collected after 6 hours and stored at −80°C. For lysate-based PBMC stimulation, the cell culture supernatants were collected after 72 hours and stored at −80°C.

### Multiplex ELISA

PBMC culture supernatants were analyzed for secreted proteins using Immune Monitoring 65-Plex Human ProcartaPlex™ Panel (Thermofisher Scientific, Carlsbad, CA), following previously documented procedures.^43,44^ This is a preconfigured multiplex immunoassay kit that measures 65 cytokines, chemokines, and growth factors for efficient immune response profiling, biomarker discovery, and validation. Tissue homogenates from all mice were evaluated using Bio-Rad 23-plex Pro Mouse Cytokine Immunoassay. Briefly, colon tissues were homogenized using Bio-Rad lysis buffer using a pestle and motorized drill for 4-6 min, centrifuged at 16,000 x g for 10 mins at 4 °C, collected supernatants, and quantified for protein concentration using Bio-Rad DC Protein Assay Kit to normalize the samples to the same total protein concentration for multiplex assay. The plates were read using the Luminex MAGPIX xMAP instrument and xPONENT® software (version 4.2, Luminex Corp, Austin, Texas, US). Standards were prepared at 1:4 dilutions (8 standards), alongside background and controls. Subsequently, concentrations of samples were determined from a standard curve using the Five Parameter Logistic (5PL) curve fit/quantification method.

### Sample preparation for Metabolomics

Anaerobic bacterial cells were treated with Nanoligomer (10 µM) designed against specific BGC for 21 hours (n= 3). Untreated bacterial cells (WT) were used as a negative control. The cells were centrifuged to remove the medium and washed once with PBS. Cell pellets were immediately extracted with ice-cold lysis/ extraction buffer (methanol: acetonitrile: water: 5:3:2). Samples were vortexed to uniformly suspend cells and incubated at 4°C for 30 min followed by centrifugation at 18000x g for 10 min. Supernatants containing the cell lysate were collected in 1.5 mL tubes and set on the SpeedVac for 2-4 hours to evaporate the extraction buffer. Dried lysates were stored at −80 °C until further use. For protein quantification, pellets were suspended in PBS by vortexing for 15 minutes at room temperature and analyzed for protein at 280 nm using Nanodrop.

### Metabolomics

Metabolomics analysis was performed at the University of Colorado School of Medicine Metabolomics Facility. Dried lysates were shipped on ice to the facility and stored at −80 °C until further use. Dried lysates were reconstituted (based on protein quantification) with 0.1% formic acid in water immediately before metabolomic analysis. Each volume added was 50% of the original dried volume. Samples were vortexed by hand at room temperature, followed by centrifugation at 4°C. All samples were analyzed twice (1 µL injection) by ultra-high-performance liquid chromatography using a Thermo Vanquish UHPLC coupled with a Thermo Q Exactive mass spectrometer in negative and positive polarity modes. For each polarized method, the UHPLC utilized a C18 column at a flow rate of 0.45 mL/min for 5 minutes. Samples entered the MS by electrospray ionization. Full technical descriptions have been published previously.^78^ Data was analyzed using Maven (1.4.20-dev-772) and quality controls were maintained as described elsewhere.^78^ Further metabolite discovery was accomplished using Compound Discoverer (3.1.0). Sample data were normalized by dividing by their respective protein quantifications and multiplying by the average quantification.

### DSS colitis mice studies

Adult female and male 6-week-old C57Bl6/J mice purchased from Jackson Laboratories were housed throughout all experiments at ∼18-23°C on a 12 hr-light/12 hr-dark cycle. Fresh water and ad-libitum food (Tekland 2918; 18% protein) were routinely provided to all cages. In week 1, all animals were fed antibiotic water (1 mg/ml each ampicillin, neomycin, and metronidazole, 0.5 mg/ml vancomycin, and 20mg/ml sugar-sweetened Kool-Aid) for 5 days, after which it was replaced with regular drinking water for 2 days. In week 2, mice were gavaged with IBD fecal slurry from 10 IBD patients (collected with approval by the Colorado Multiple Institutional Review Board protocol #17-0977), and allowed 1-week for stable colonization. In week 3, mice were provided DSS in their drinking water (3% DSS by weight for 5 days in the acute model). After 5 days, DSS water was removed and replaced with normal drinking water.

For the acute DSS model, the mice were euthanized on day 7, and colon tissue was extracted. For the DSS chronic model, the mice were provided with 2.5% DSS (wt/vol) in the drinking water for 5 days, then 5 days on regular drinking water, and this cycle was repeated 3 times. At the end of 3 cycles, the mice were euthanized, and colon extracted.

For respective IP treatments, the mice were given 150 mg/kg dose injections 3x per week, for the duration of study post DSS induction. For oral treatments, the mice were given size M oral capsule using the applicator (Torpac).

The animal weights and stool were monitored daily while on DSS water. Animals were consistently health-checked by the veterinary staff.

### TNF***^Δ^***^ARE/+^ mouse model

TNF^ΔARE/+^ transgenic mice were bred at the CU Anschutz vivarium. Littermate TNF^ΔARE/+^ mice were used for vehicle and Nanoligomer treatments (as described above), while TNF-negative mice (TNF^+/+^) were used for negative controls. Similar protocols for inducing gut dysbiosis were followed where 4-week-old mice were treated with antibiotic water for 5 days and then gavaged using fecal slurry and microbial spike. After one week, the mice were treated for 4-weeks and monitored for colitis. At the end of the 4 weeks, the 10-week-old mice were euthanized, their colons harvested, and analyzed using multiplexed ELISA.

### Oral formulation

Size M (mice) oral capsules (Torpac) were completely filled with dried Nanoligomer NI112 mixed with dicalcium phosphate (LFA) in a 50:50 ratio. The weighed capsules (∼3mg/capsule) had 75mg/kg API dosing for 20 mg mice. The capsules were then coated with Eudragit (Evonik) enteric polymer, dried, and given to mice using an oral gavage applicator (Torpac).

### Tissue collection

After deep anesthetizing with isoflurane, ∼1 ml of blood was removed via cardiac puncture followed by cervical dislocation. The colons were dissected and either flash-frozen (for ELISA) or preserved in 10% normal buffered formalin (NBF) using the Swiss-roll technique. The flash-frozen samples were frozen on dry ice and stored at −80°C until further processing.

### Hematoxylin and Eosin (H&E) staining for tissue morphology

Colons were fixed in 10% neutral buffered formalin at room temperature for at least 48 hours. Tissues were processed using a Leica TP1020 Automatic Benchtop Tissue Processor and embedded in paraffin wax (Cancer Diagnostics, Cat #: EEPAR56). Tissues were sectioned and mounted on positively charged glass slides (Superfrost Plus, Cancer 232 Diagnostics, Cat #: 4951) for staining and analysis. One section per animal was deparaffinized and stained with hematoxylin (Cancer Diagnostics, Cat#: #HTV-4) and eosin (Cancer Diagnostics, Cat#: #ETV) (H&E) for the determination of histopathological changes. Whole tissue images were taken for analysis.

### Statistics

The figure legends specify the statistical tests conducted, the number of mice involved, and the reported p-values. To evaluate variances among groups, a one-way ANOVA. Microsoft Excel and GraphPad Prism were employed for data analysis, and GraphPad Prism software was used for data presentation.

## ASSOCIATED CONTENT

### Supporting Information

Supporting Figure S1 shows all anaerobic strains used in the study, and figures S2-S8, with 65-plex Multiplexed human ELISA plots for *in vitro* screening, in the gut-immune model.

### Author Information

#### Corresponding Authors

*, *

#### Author Contributions

P.N. conceived the idea, designed the experiments, and synthesized the Nanoligomers. S.S. and A.C. designed the *in vitro* culture studies. S.S. cultured all microbial strains, conducted Nanoligomer treatments, qPCR, and metabolomic studies. S.S. and V.G. conducted all the PBMC *in vitro* experiments and biochemical characterization for *in vitro* and *in vivo* studies. K.A.K. designed all animal studies. C.L.L. and K.A.K. conducted the DSS mouse and TNF^ΔARE/+^ mouse studies and histology. S.S., A.C., and P.N. conducted the ELISA measurements. P.N. and A.C. wrote the manuscript with input from all the authors. All authors read the manuscript and provided input.

## ACKNOWLEDGMENTS

Authors acknowledge financial support from NIH SBIR Award R43DK132980 (NIDDK).

## SUPPORTING INFORMATION

### Supplementary Figures

**Fig. S1.**
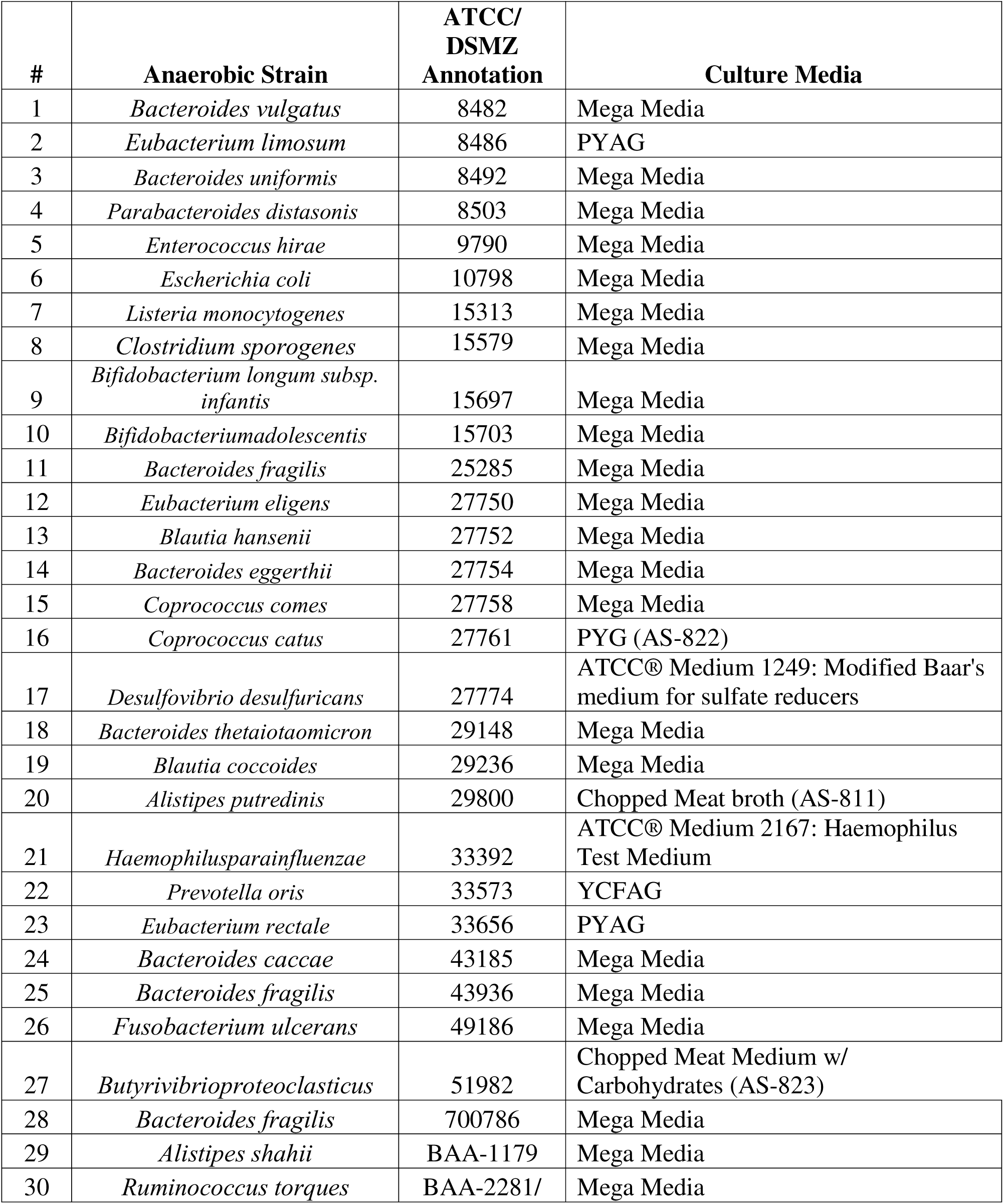
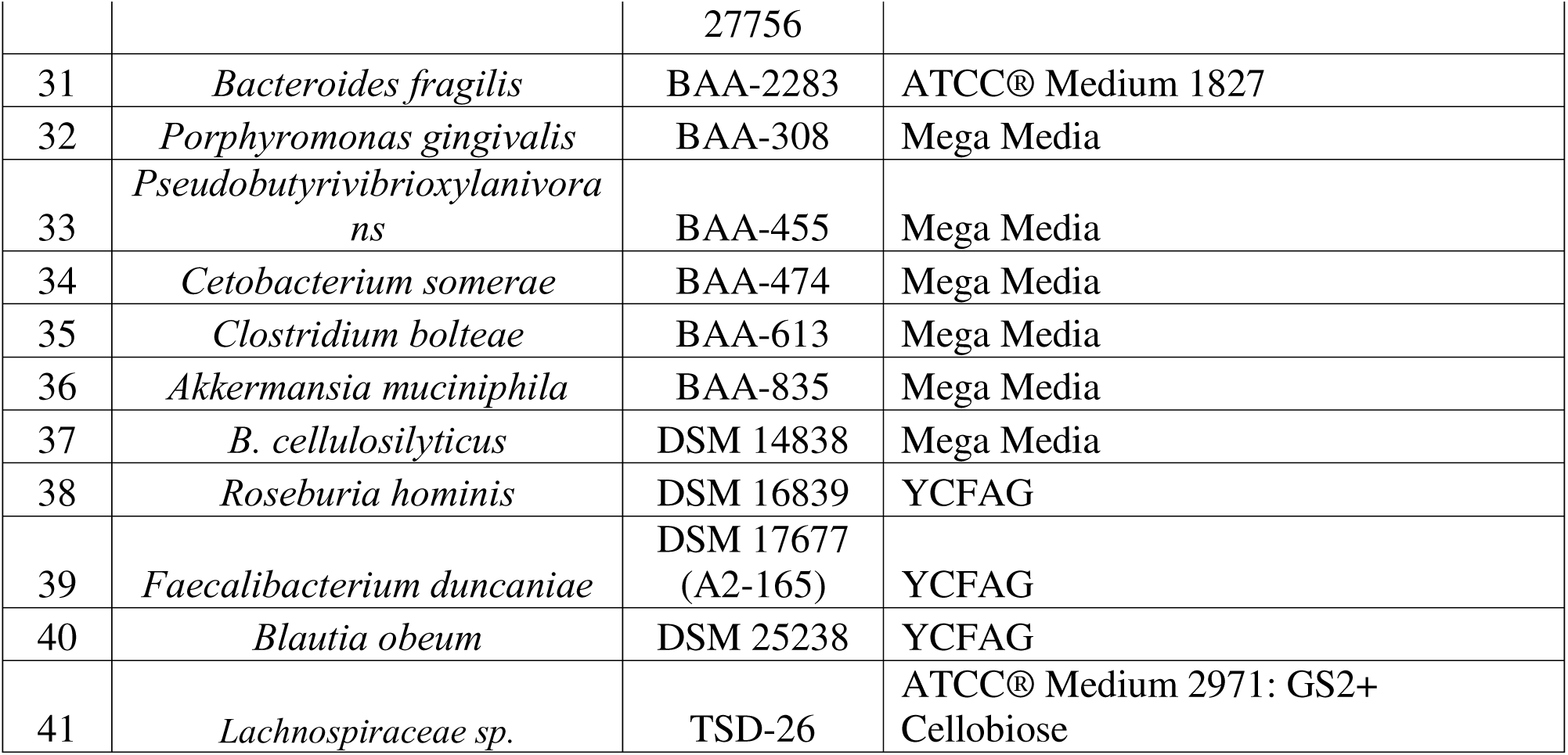
Cultured gut anaerobe species used in this study for microbiome-targeted IBD therapy. PYAG: Peptone-yeast extract (PY) broth supplemented with 1% glucose and 33 mM sodium acetate medium; YCFAG : Yeast extract, casitone, fatty acid, and glucose medium; PYG: Peptone-Yeast extract supplemented with 1% glucose

**Fig. S2.**
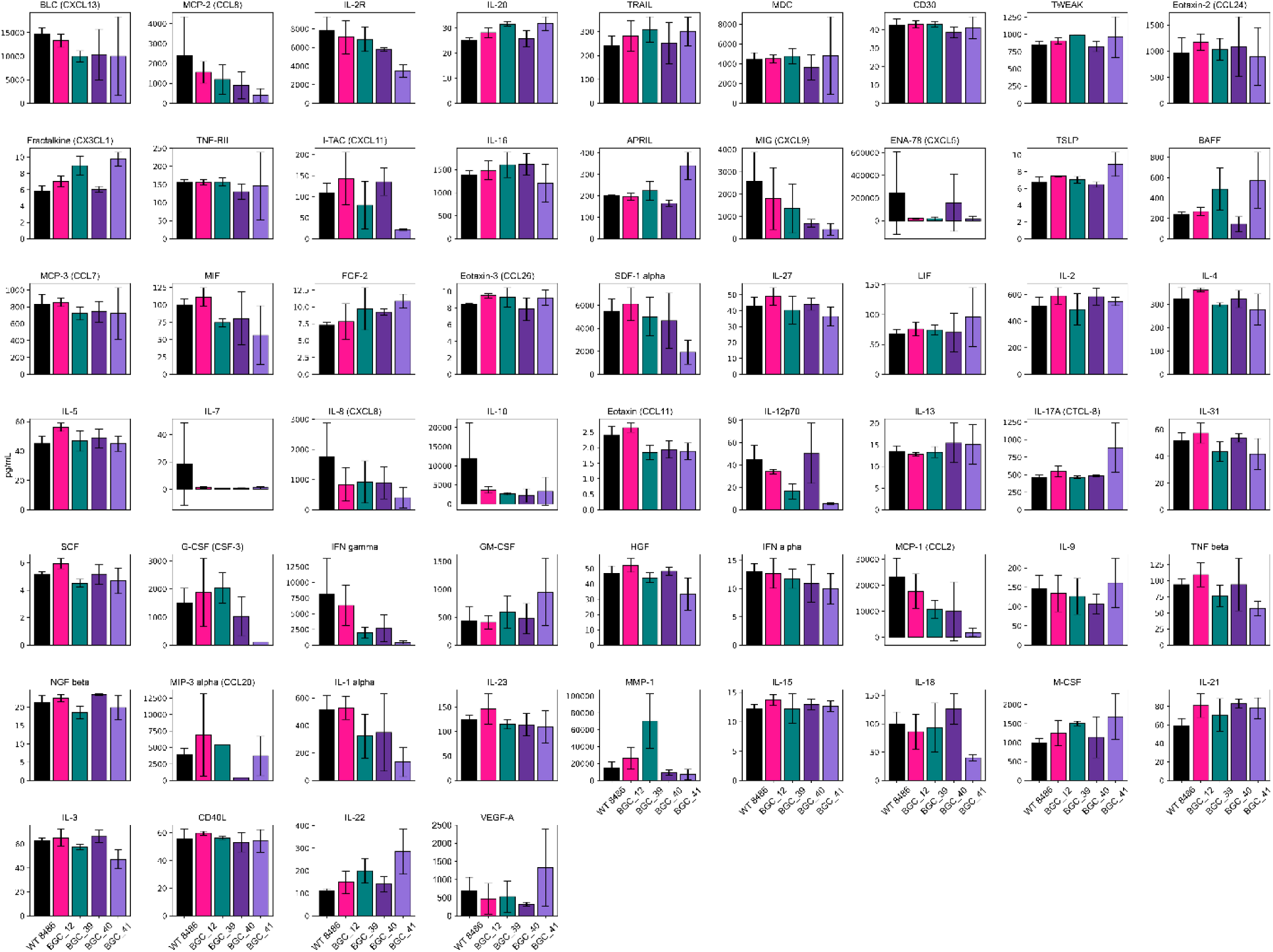
Screening microbiome targets in *Eubacterium limosum* (8486) using 65-plex multiplexed ELISA analysis of human cytokines and chemokines. Donor-derived human PBMCs were treated using WT cell culture lysates (normalized for cell count using optical density OD) of *E. limosum* (8486), and 4 different annotated biosynthetic genes (BGCs 12, 39, 40, and 41) suppressed using respective Nanoligomer targeting. The PBMC supernatants were subsequently analyzed using 65 different human cytokines and chemokines, presented here, for assessing the impact of different microbiome-targeted treatments.

**Fig. S3.**
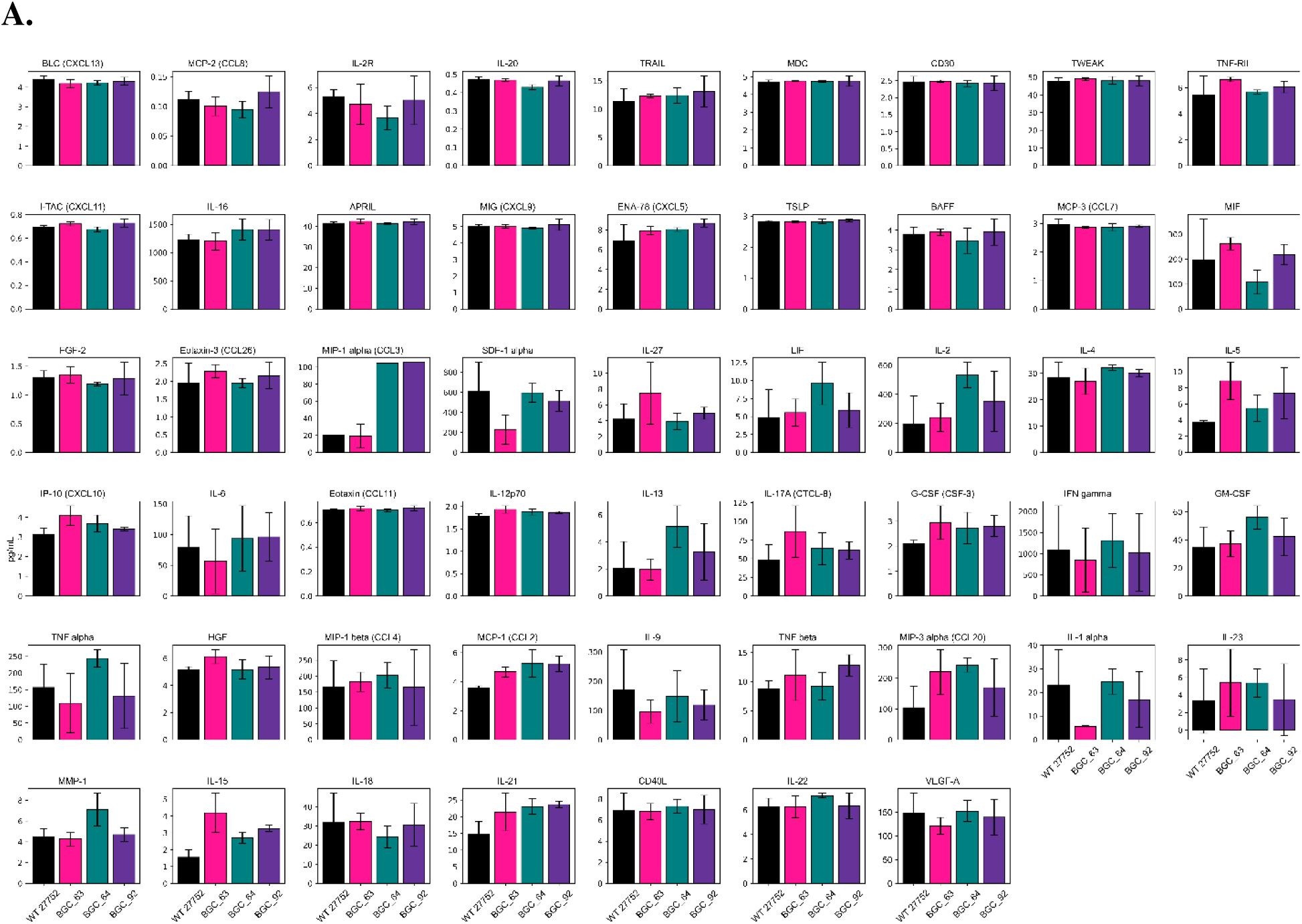
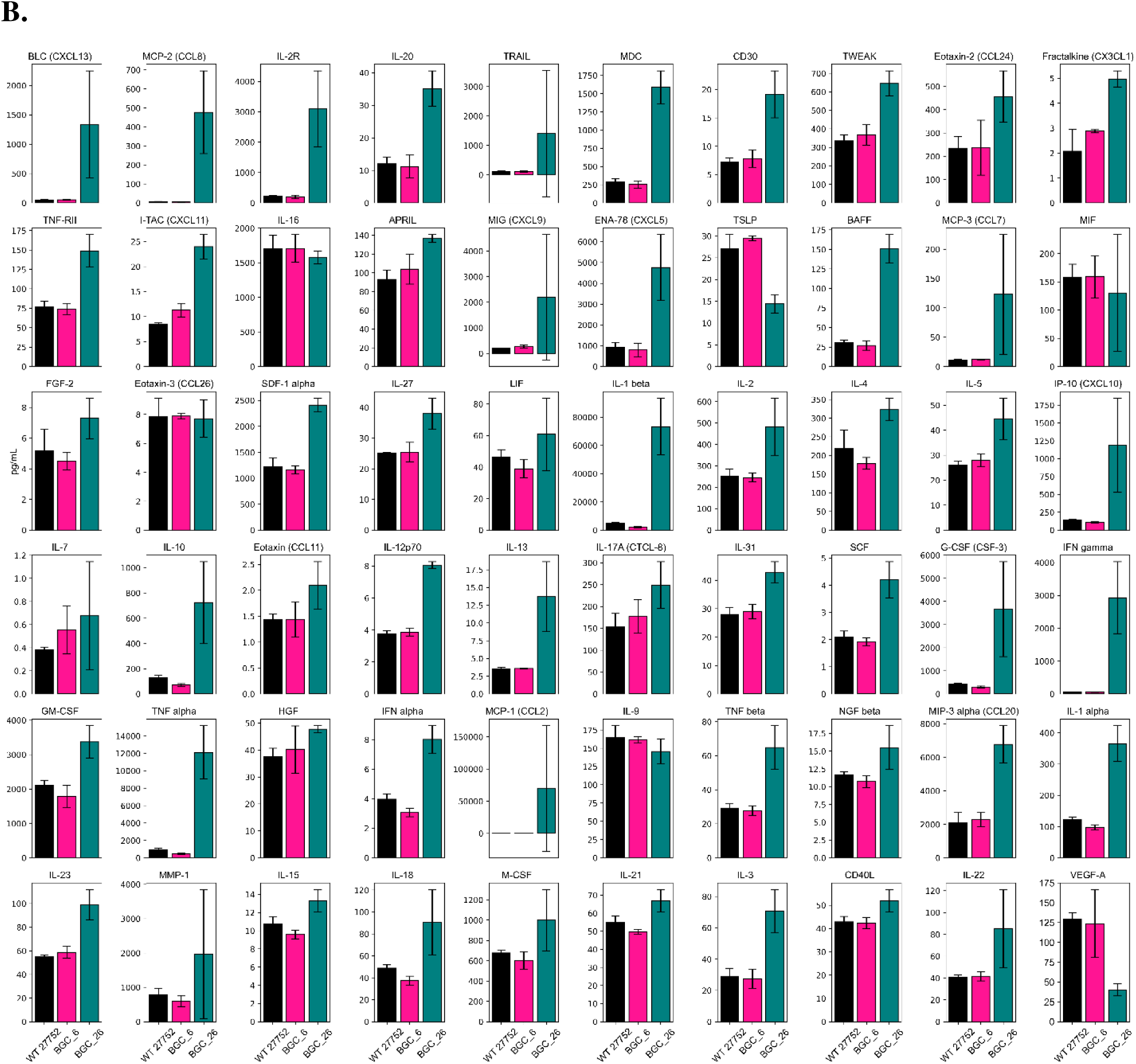
Screening microbiome targets in *Blautia hansenii* (27752) using 65-plex multiplexed ELISA analysis of human cytokines and chemokines. Donor-derived human PBMCs were treated using WT cell culture lysates (normalized for cell count using optical density OD) of *B. hansenii* (27752), and 5 different annotated biosynthetic genes, in **A**. BGCs 63, 64, 92, and **B**. BGCs 6, 26 were suppressed using respective Nanoligomer targeting. The PBMC supernatants were subsequently analyzed using 65 different human cytokines and chemokines, presented here, for assessing the impact of different microbiome-targeted treatments.

**Fig. S4.**
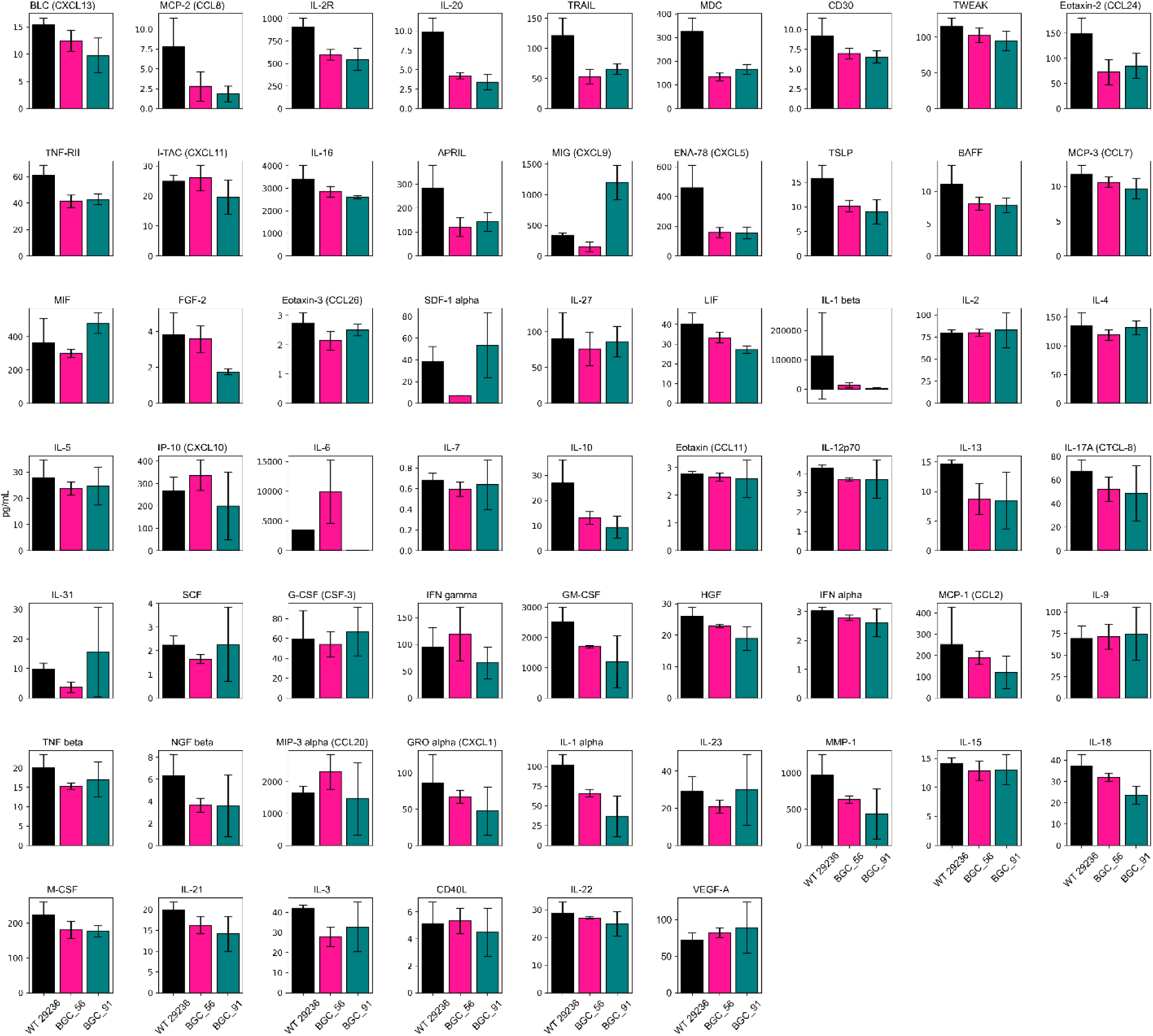
Screening microbiome targets in *Blautia coccoides* (29236) using 65-plex multiplexed ELISA analysis of human cytokines and chemokines. Donor-derived human PBMCs were treated using WT cell culture lysates (normalized for cell count using optical density OD) of *B. coccoides* (29236), and 2 different annotated biosynthetic genes (BGCs 56 and 91) suppressed using respective Nanoligomer targeting. The PBMC supernatants were subsequently analyzed using 65 different human cytokines and chemokines, presented here, for assessing the impact of different microbiome-targeted treatments.

**Fig. S5.**
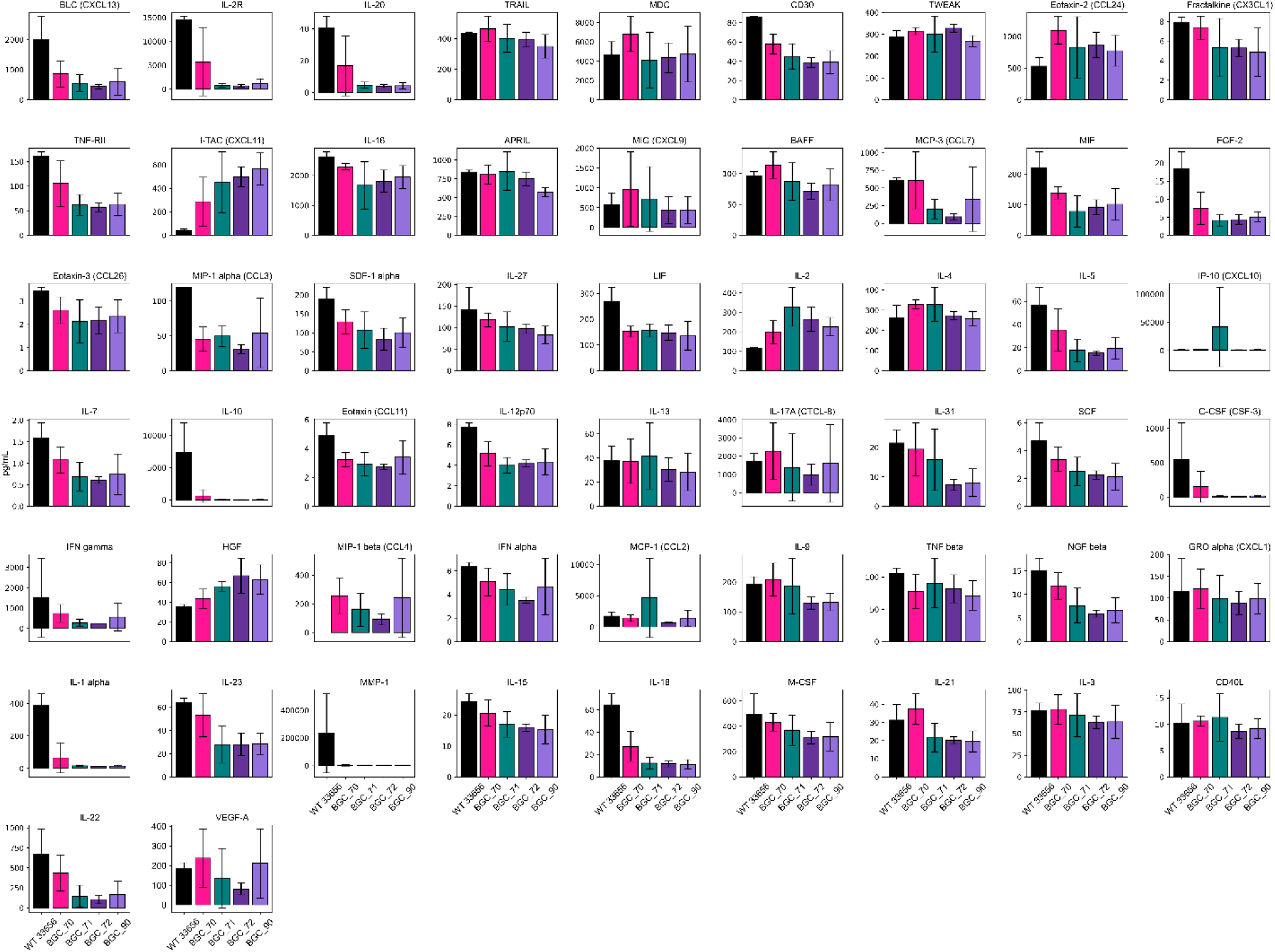
Screening microbiome targets in *Eubacterium rectale* (33656) using 65-plex multiplexed ELISA analysis of human cytokines and chemokines. Donor-derived human PBMCs were treated using WT cell culture lysates (normalized for cell count using optical density OD) of *E. rectale* (33656), and 4 different annotated biosynthetic genes (BGCs 70, 71, 72, and 90) suppressed using respective Nanoligomer targeting. The PBMC supernatants were subsequently analyzed using 65 different human cytokines and chemokines, presented here, for assessing the impact of different microbiome-targeted treatments.

**Fig. S6.**
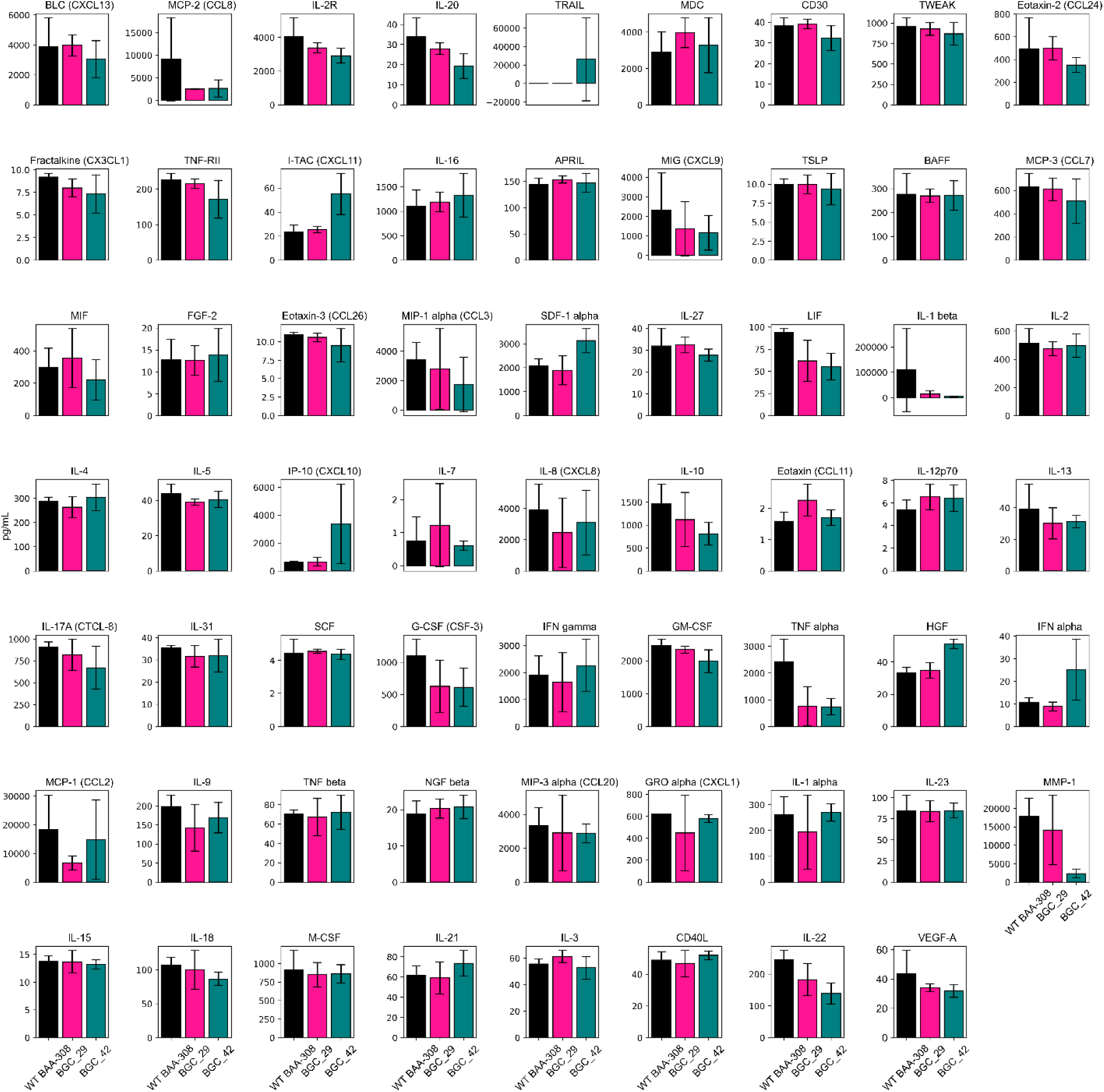
Screening microbiome targets in *Porphyromonas gingivalis* (BAA-308) using 65-plex multiplexed ELISA analysis of human cytokines and chemokines. Donor-derived human PBMCs were treated using WT cell culture lysates (normalized for cell count using optical density OD) of *P. gingivalis* (BAA-308), and 2 different annotated biosynthetic genes (BGCs 29, and 42) suppressed using respective Nanoligomer targeting. The PBMC supernatants were subsequently analyzed using 65 different human cytokines and chemokines, presented here, for assessing the impact of different microbiome-targeted treatments.

**Fig. S7.**
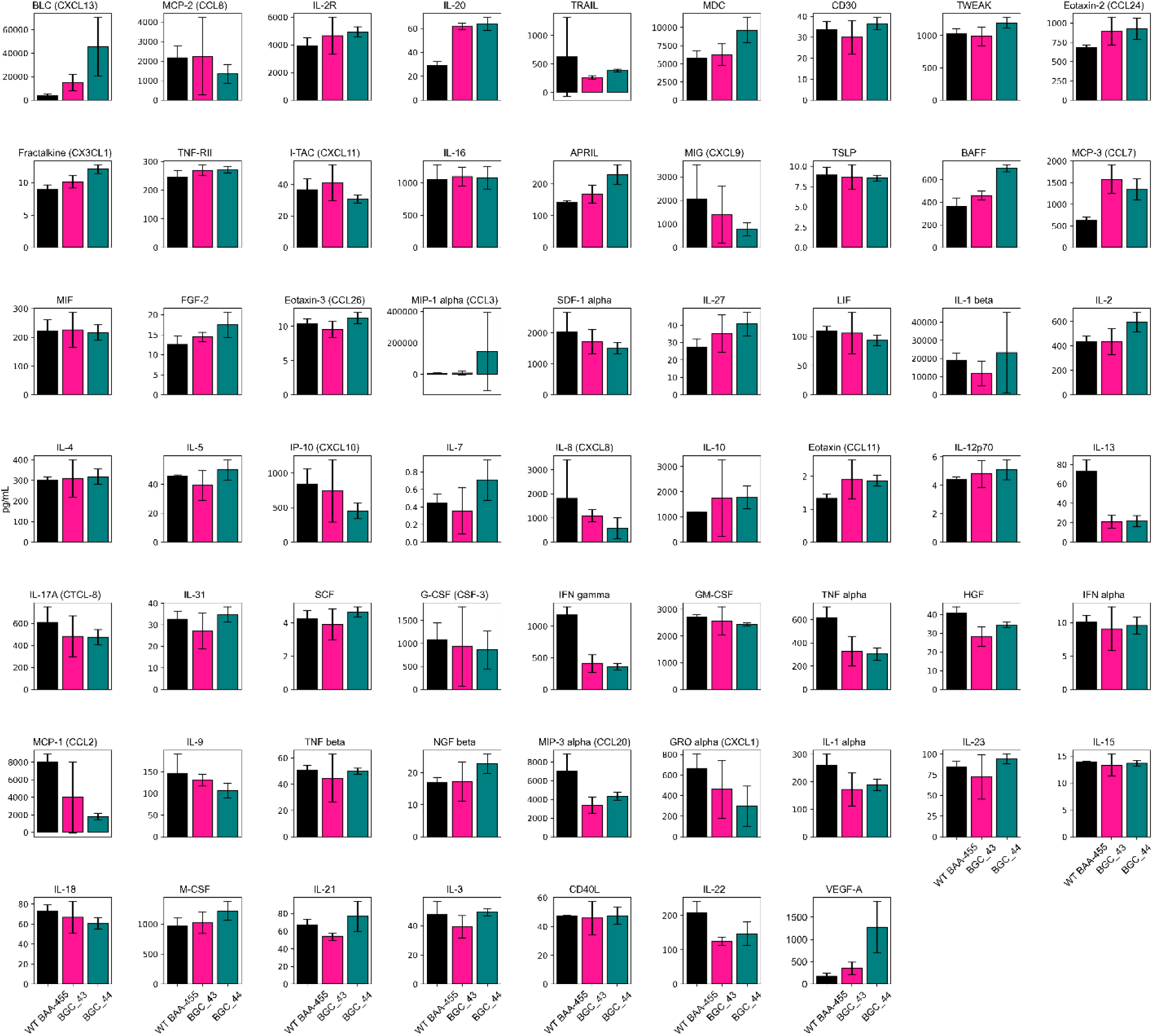
Screening microbiome targets in *Pseudobutyrivibrio xylanivorans* (BAA-455) using 65-plex multiplexed ELISA analysis of human cytokines and chemokines. Donor-derived human PBMCs were treated using WT cell culture lysates (normalized for cell count using optical density OD) of *P. xylanivorans* (BAA-455), and 2 different annotated biosynthetic gene (BGCs 43, and 44) suppressed using respective Nanoligomer targeting. The PBMC supernatant were subsequently analyzed using 65 different human cytokines and chemokines, presented here, for assessing the impact of different microbiome-targeted treatments.

**Fig. S8.**
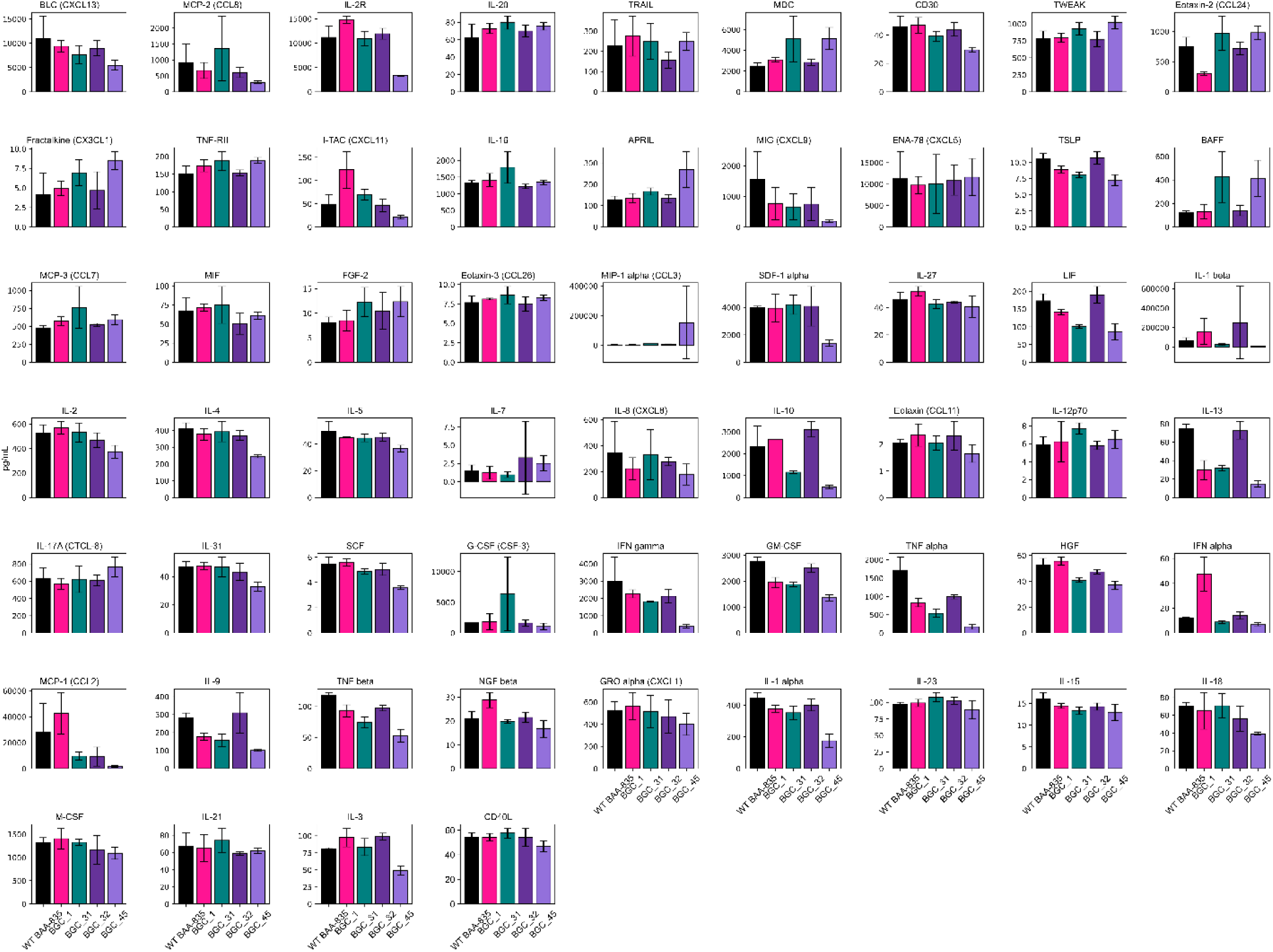
Screening microbiome targets in *Akkermansia muciniphila* (BAA-835) using 65-plex multiplexed ELISA analysis of human cytokines and chemokines. Donor-derived human PBMCs were treated using WT cell culture lysates (normalized for cell count using optical density OD) of *A. muciniphila* (BAA-835), and 4 different annotated biosynthetic genes (BGCs 1, 31, 32, and 45) suppressed using respective Nanoligomer targeting. The PBMC supernatants were subsequently analyzed using 65 different human cytokines and chemokines, presented here, for assessing the impact of different microbiome-targeted treatments.

